# Ripply1 and Gsc collectively suppress anterior endoderm differentiation from prechordal plate progenitors

**DOI:** 10.1101/2023.05.29.542713

**Authors:** Tao Cheng, Xiang Liu, Yang Dong, Yi-Meng Tian, Yan-Yi Xing, Chen-Yi Chen, Cong Liu, Yun-Fei Li, Ying Huang, Ding-Hao Zhuo, Xiao Xu, Jing-Yun Luan, Xin-Xin Fu, Zi-Xin Jin, Jing Mo, Xiang Xu, Hong-Qing Liang, Peng-Fei Xu

## Abstract

During gastrulation, the mesendoderm is firstly specified by morphogens such as Nodal, and then segregates into endoderm and mesoderm in a Nodal concentration-dependent manner. However, the mechanism underlying the segregation and crosstalk of different sub-groups within the meso- and endoderm lineages remains unclear. Here, taking zebrafish prechordal plate (PP) and anterior endoderm (Endo) as research model, using single-cell multi-omics and live imaging analyses, we show that anterior Endo progenitors originate directly from PP progenitors. A single-cell transcriptomic trajectory analysis of wild-type, *ndr1* knockdown and *lft1* knockout Nodal explants confirms the diversification of anterior Endo fate from PP progenitors. Gene Ontology (GO) enrichment analysis identifies that the change of chromatin organization potentiates the segregation of anterior endodermal cell fate from PP progenitors. Single-cell ATAC & RNA sequencing further reveals that two transcriptional regulators, *gsc* and *ripply1*, exhibit varied activation patterns in PP and anterior Endo cell trajectories at both the chromatin and RNA expression levels. We further demonstrate that Ripply1 functions coordinately with Gsc to repress anterior endodermal cell fate by directly binding to the *cis*-elements of *sox32*. Modulating the expression levels of these regulators tilts the cell fate decision between the PP and anterior Endo.

## Introduction

During gastrulation, instructed by genetic and epigenetic signals, naïve cells from blastula progressively acquire distinct cell fates and are allocated to specific domains within the embryo^1,2^. Nodal, a member of the transforming growth factor β (TGF-β) superfamily, acts as a morphogen^1,3–5^ and plays highly conserved roles in mesendoderm induction and patterning^6–8^.

It is widely accepted that Nodal induces endodermal and mesodermal cell fates via its concentration gradient^8–10^. Higher Nodal signaling activity bias cells toward endoderm, while lower levels are thought to promote mesoderm induction^11,12^. Mesodermal and endodermal common progenitors are induced by Nodal and mixed together at the onset of gastrulation^1^, while it remains elusive how the common progenitors of mesendoderm took on distinct lineages. Several recent studies reported that multiple important signaling pathways interacted with Nodal to drive mesendoderm separation^2,13–15^. Notably, lateral endoderm specification may comply with a stochastic cell fate switching model in zebrafish^16^.

The specification of dorsal mesendoderm requires the highest level of Nodal signaling, as evidenced by that this cell population was first to be depleted when Nodal signaling was gradually inhibited^11^. Both the PP and anterior Endo originate from anterior mesendoderm (derived from the dorsal mesendoderm), and the commitment of these cell fates is regulated by similar levels of Nodal signaling. However, the mechanisms governing the separation of cell fates within the anterior mesendoderm remain unclear. Previous studies show that Nodal signaling enhances the duration of cell-cell contact in PP cells, creating a positive feedback loop between cell-cell contact duration and cell fate specification^17^. However, what factors downstream of Nodal pathway play major roles in segregating these two cell fates, when and how PP and anterior Endo cells can be distinguished transcriptionally are the questions remaining to be investigated.

Previous research has shown that Gsc, whose expression can be induced by Nodal, acts as a repressor to suppress anterior endoderm specification by binding to the promoter region of *sox17*^18^. However, investigations into loss-of-function of Gsc in zebrafish have not yielded conclusive evidence supporting its significant roles in mesoderm and/or endoderm specification^19,20^. These results imply that there might be additional, unidentified transcriptional repressors that work in collaboration with Gsc and possess a redundant function in suppressing the specification of anterior endoderm.

In this study, we constructed a single-cell transcriptional trajectory to delineate the cell state segregation between PP and anterior Endo in zebrafish embryos and Nodal-injected zebrafish explants^21,22^. Single-cell transcriptional trajectory and live imaging analyses demonstrated that anterior Endo cells originated and were specified from PP progenitors. Interestingly, this cell trajectory separation can be traced back to as early as the onset of gastrulation. To achieve a more comprehensive understanding, we integrated single-cell datasets from wild-type, *ndr1* (Nodal ligand) knockdown and *lft1* (Nodal inhibitor) knockout Nodal explants at shield stage to construct an extended single-cell trajectory, which fully captured the branching event between PP and anterior Endo fates and unveiled a potential involvement of epigenetic regulation in this process. This finding was further supported by the experimental evidence that perturbation of the chromatin remodeler *srcap* could affect the cell specification of PP and anterior Endo. Furthermore, combining the multi-omic analysis and gain-/loss-of-function experiments, we found that a transcriptional repressor, Ripply1, together with Gsc, collectively suppresses anterior endoderm specification by directly binding to the regulatory elements of *sox32*.

## Results

### Anterior Endo originates from PP progenitors in zebrafish

Both anterior Endo and PP are derived from embryonic shield and require high Nodal signaling to be specified^11,18,23^. To investigate how these two cell lineages are further segregated, we collected the single-cell RNA-seq datasets from the published studies^24,25^, identified the subpopulations representing the anterior Endo and PP cells, and reconstructed their trajectory branching tree (Figures 1A-1C and S1A). We found that these two trajectories started to be separated at shield stage, and their common progenitors highly expressed PP marker genes, like *gsc, frzb*, etc. (Figures 1C and S1A). Then, we further explored the dynamic molecular signatures of these two trajectories during their fate determination (Figures S1B-S1E). Pseudotime analysis revealed that PP and anterior Endo exhibited distinct differentiation states by the end of gastrulation (Figure S1B). Of note, compared to *sox32*, *gsc* displayed earlier transcriptional differences in these two cell trajectories (Figure S1C). The cell trajectory prediction analysis^26^ also revealed that the progenitor cells resembled more of the PP fate (Figures 1D, 1E and S1F), and similar result was obtained using another single-cell RNA-seq dataset (Figures S1G-S1I). Therefore, our analysis suggest that anterior Endo and PP originate from the same progenitor with transcriptome profile more similar to PP fate. Anterior Endo becomes transcriptionally distinguishable from PP as early as at the shield stage.

**Figure 1.**
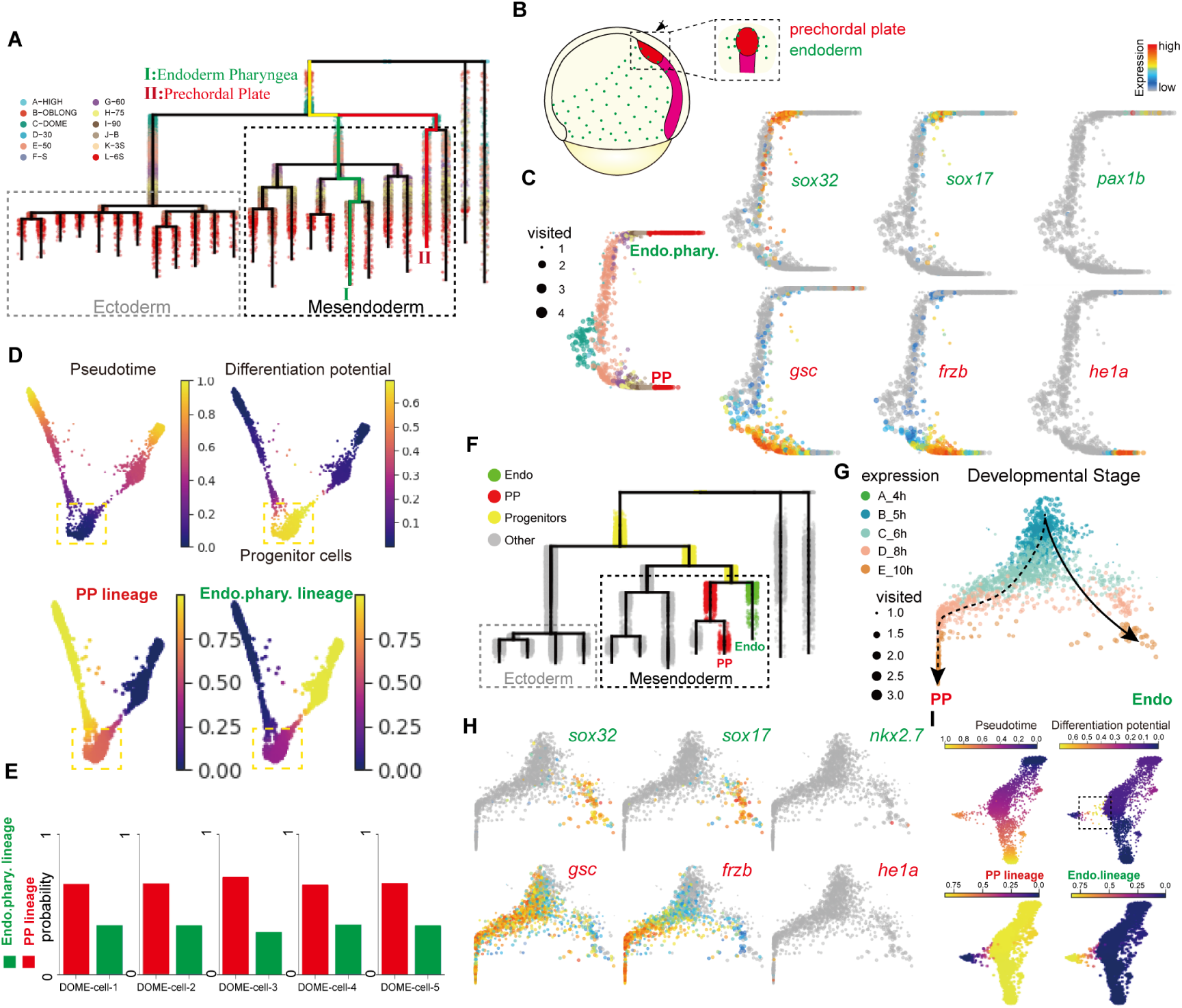
Single-cell trajectory analysis for studying PP and anterior Endo specification. (A) URD differentiation tree of zebrafish embryos from blastula stage to somitogenesis stage^73^. Cells are colored according to developmental stages. Gray dotted frame indicates the ectoderm trajectories and black dotted frame indicates the mesendoderm trajectories. Green line shows the pharyngeal endoderm trajectory (anterior endoderm, roman number I), red line shows the prechordal plate trajectory (roman number II) and yellow line indicates their progenitors. (B) Simplified schematic showing the patterns of anterior endoderm and axial mesoderm of zebrafish embryos at 75% epiboly. Dotted frame indicates the enlarged view of dorsal anterior region, showing that anterior Endo sporadically locates near PP. (C) URD branchpoint plots of the PP and anterior Endo development from scRNA-seq data of zebrafish embryos^73^, showing pseudotime (x-axis) and random walk visitation preference from pharyngeal endoderm to prechordal plate domains (y-axis). Cells are colored by developmental stages (left) and expression of anterior Endo markers (*sox32*, *sox17* and *pax1b*; right top) and PP markers (*gsc*, *frzb* and *he1a*; right bottom). (D) Force-directed layout of PP and anterior Endo cells of zebrafish embryos from blastula stage to somitogenesis stage. Cells are colored by Palantir^26^ pseudotime (top left), differentiation potential (top right) and branch probabilities of PP (bottom left) and anterior Endo (bottom right). (E) Branch probabilities of five cells randomly selected from progenitor cells. Bars are colored by cell types. (F) URD differentiation tree of Nodal explants from blastula stage to the end of gastrulation. Gray dotted frame indicates the ectoderm trajectories and black dotted frame indicates the mesendoderm trajectories. The green, red and yellow dots indicate the Endo cells (anterior Endo), PP cells and the progenitor cells respectively, and the gray dots indicate other cells. (G-H) URD branchpoint plots of the PP and anterior Endo development from scRNA-seq data of Nodal explants showing pseudotime (y-axis) and random walk visitation preference from PP to anterior Endo domains (x-axis). Cells are colored by developmental stages (G) and expression of anterior Endo markers (*sox32*, *sox17* and *nkx2.7*; top) and PP markers (*gsc*, *frzb* and *he1a*; bottom) (H). (I) Force-directed layout of PP and anterior Endo cells of Nodal explants from blastula stage to the end of gastrulation. Cells are colored by the levels of Palantir^26^ pseudotime (top left), differentiation potential (top right) and branch probabilities of PP (bottom left) and anterior Endo (bottom right).

Cells from hatching gland (derived from PP) and pharyngeal pouch (derived from anterior Endo) are distant on the transcriptome-based phylogenetic tree of zebrafish embryos^25^. Thus, the resolution of this dataset may not be sufficient to identify the exact trajectory separation point between PP and anterior Endo. To investigate the key molecules that regulate the PP and anterior Endo separation, we employed our previously published Nodal induced embryonic explant system (referred to Nodal explant hereafter), which encompasses the axial mesendoderm, along with the PP and anterior Endo fates^22^. Notably, the single-cell trajectory tree of Nodal explants showed a closer transcriptomic relationship between the PP and anterior Endo (Figure 1F). Thus, we reconciled the single-cell transcriptome dynamics to look for the branching point of PP and anterior Endo in the Nodal explants (Figures 1G, 1H, S2A and S2B). Consistent with the observations in wild-type embryos, the common progenitors of these two cell trajectories highly expressed the PP marker gene (*gsc*), and the segregation between PP and anterior Endo had already occurred by 6 hpf (hours post fertilization) (Figures 1G, 1H and S2B). Cell trajectory prediction analysis on the Nodal explant datasets further indicated that the common progenitor cells were transcriptionally similar to PP (Figures 1I and S2I), which aligned with our previous observations in wild-type embryos (Figures 1D, 1E, S1H and S1I).

To explore the molecular signatures during the specification of PP and anterior Endo, we performed differential expression analysis and the gene module identification analysis for these two cell trajectories in both wild-type embryos and Nodal explants^27^. These genes enriched GO terms^28^ were highly correlated with the developmental processes and functions of these two cell types (Figures S2C-S2H).

To further validate this observation and investigate the dynamics of cell fate segregation between anterior Endo and PP during development, we performed live imaging analysis on embryos of Tg(*gsc*:EGFP;*sox17*:DsRed) (Figures 2A and 2B). Notably, we observed a gradual emergence of *sox17-*positive cells from a subset of the *gsc*-positive cells (Figures 2A-2D, S3A-S3D and Movies S2, S3). And this cell fate transition was also evident in Nodal explants constructed from Tg(*gsc*:EGFP;*sox17*:DsRed) embryos (Figures S3E-S3H and Movie S1). Furthermore, we generated *sox17*:Cre and *gsc*:loxP-STOP-loxP-mCherry constructs (Figure S4D) and co-injected them with transposase mRNA into one-cell stage embryos (Figure S4A). Interestingly, distinct mCherry-positive cells were observed in the injected embryos, providing clear evidence that *gsc*-positive cells can give rise to *sox17*-positive cells (Figures S4B and S4C).

**Figure 2.**
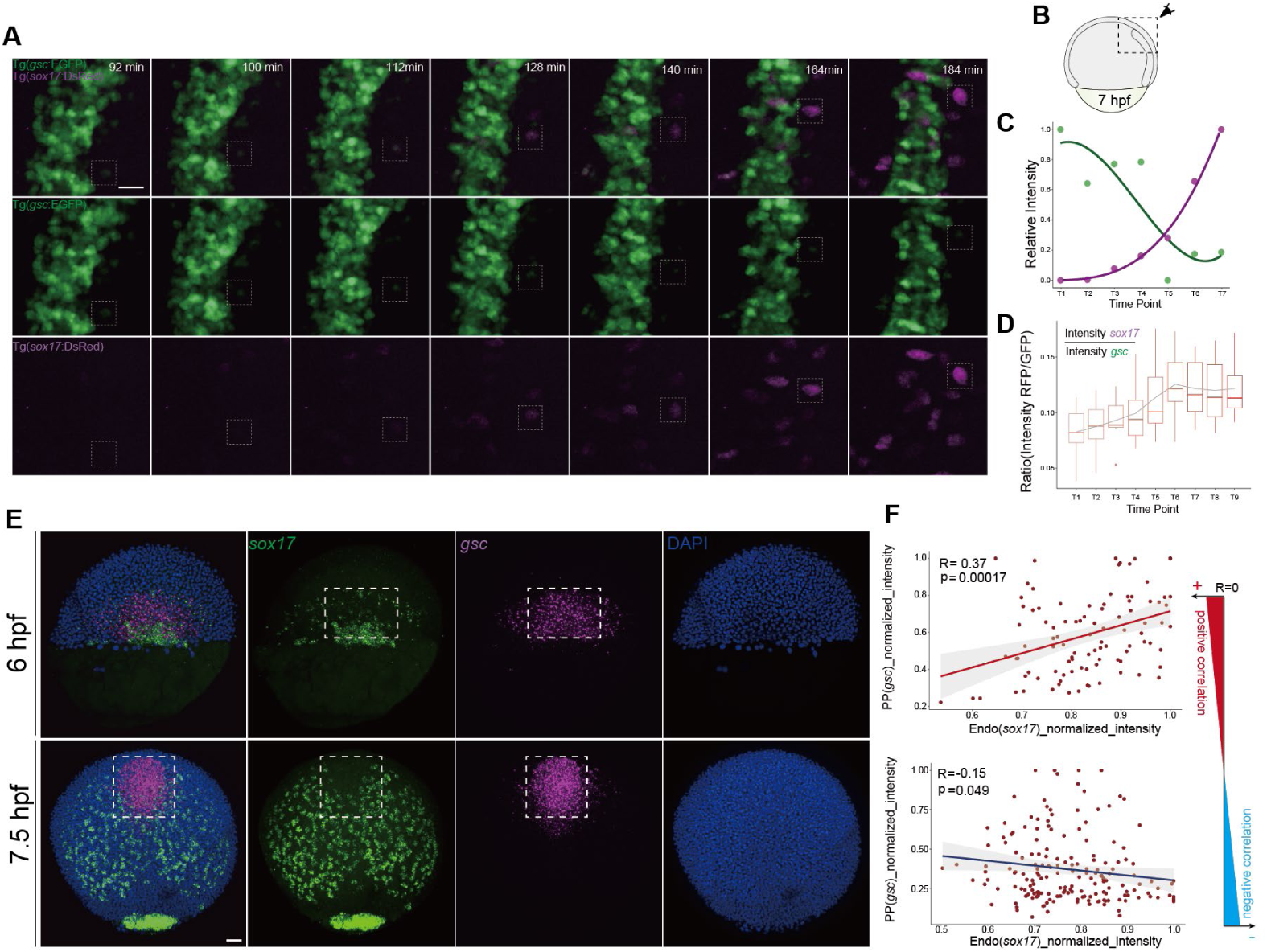
Live imaging analyses for Tg(*gsc*:EGFP;*sox17*:DsRed) embryos. (A) Time series of 3D reconstruction of a representative Tg(*gsc*:EGFP;*sox17*:DsRed) embryo. (B) Schematic diagram indicating the focused region in the Tg(*gsc*:EGFP;*sox17*:DsRed) embryo. (C) Temporal profiles of relative intensity of *sox17* (RFP) & *gsc* (GFP) in the highlighted cell from (A). (D) Temporal profiles of the ratio between RFP intensity and GFP intensity in multiple tracked cells. (E) Double-color RNA-fluorescence *in situ* hybridization (FISH) of *sox17* and *gsc* in wild-type embryos at 6 hpf (top) and 7.5 hpf (bottom). (n = 5/5, 6 hpf; n = 5/5, 7.5 hpf; n: embryos were imaged, expression observed/total imaged) (F) Scatter plots and inferred linear regression comparing the relative intensities of *sox17* and *gsc* in wild-type embryos at 6 hpf (top) and 7.5 hpf (bottom). Each FISH experiment was performed for at least 3 independent replicates (technical replicates); more than 30 embryos were analyzed. Scale bar: 30 µm (A) and 50 µm (E).

We co-stained *gsc* and *sox17* in embryos and in Nodal explants respectively at different time points during gastrulation (Figures 2E and S3I). By assessing the expression levels of *sox17* and *gsc* in *sox17*-positive anterior mesendoderm cells, we observed a positive correlation between *sox17* and *gsc* expression at the onset of gastrulation. However, with advancing development, *sox17* and *gsc* expression exhibited a negative correlation in both embryos and Nodal explants (Figures 2F and S3J). This suggests that the increase of *sox17* expression during anterior Endo differentiation is accompanied by the simultaneous reduction of *gsc* expression. These findings align with our live imaging experiments (Figures 2A, S3B and S3F). In summary, combing single-cell transcriptomic trajectory analyses and live imaging experiments using embryos and explants, we demonstrated that anterior Endo cells were derived from PP progenitor cells in zebrafish.

### Nodal-Lefty regulatory loop is needed for PP and anterior Endo fate specification

In zebrafish, the PP and anterior Endo lineages are physically in close proximity and direct contact^17^. During development, interactions through cell-cell contacts between neighboring cell types often play a crucial role in promoting cell fate segregation^17,29^. We hypothesized that communications between PP and anterior Endo cells may influence each other’s cell fate specification. We employed LIANA^30^, a ligand-receptor analysis framework, to investigate the molecular signals involved in these interactions using single-cell RNA-seq datasets. Our analysis revealed strong interactions between PP and anterior Endo cells in both wild-type embryos and Nodal explants (Figures S5C-S5F). Interestingly, key molecular factors in the Nodal-Lefty regulatory loops (Figure 3D), such as *ndr1*, *lft1* and *lft2*, were significantly enriched in the interaction between PP and anterior Endo cells (Figures 3A, 3B, S5A, S5B and S5G-S5I).

**Figure 3.**
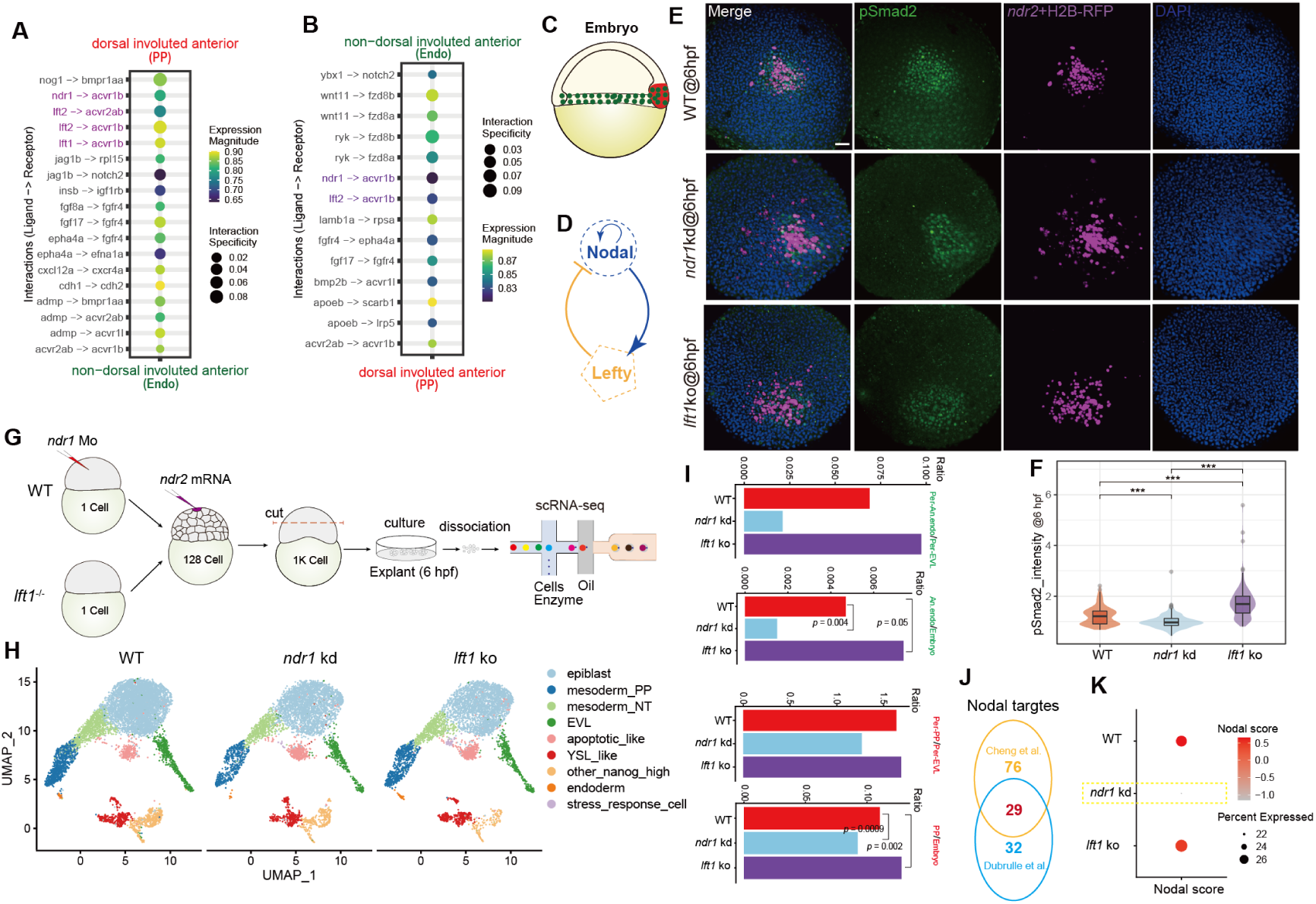
Nodal-Lefty regulatory loops are involved in cell fate separation between anterior Endo and PP. (A and B) Dot plots visualizing ligand-receptor interactions between PP and anterior Endo. The interactions from PP (ligand) to anterior Endo (receptor) and from anterior Endo (ligand) to PP (receptor) are plotted respectively in A and B. (C) Schematic diagram illustrating PP (red) and anterior Endo (green) in zebrafish embryos at 6 hpf. (D) Simplified schematic showing the interaction network between Nodal and Lefty. (E) 3D reconstruction of representative confocal-scanned images showing pSmad2 levels in zebrafish embryonic animal pole stimulated by Nodal injection at 128-cell stage (WT, n = 3/4; *ndr1*kd, n = 3/4; *lft1*ko, n = 4/4; n: embryos were imaged, observed/total imaged). (F) Comparisons of pSmad2 levels in wild-type embryos, *lft1* mutants and *ndr1* morphants at 6 hpf (quantified from E). Statistical differences between two samples were evaluated by Student’s t-test. *indicates P-value < 0.05, **indicates P-value < 0.01 and ***indicates P-value < 0.001. (G) Schematic diagram showing the experimental workflow of single-cell RNA sequencing of Nodal explants constructed from *lft1* mutants and *ndr1* morphants at 6 hpf. (H) Overall UMAP plot of integrated single-cell datasets of Nodal explants generated from wild-type embryos (left), *ndr1* morphants (middle) and *lft1* mutants (right). (I) Histograms showing the ratios of the cell proportions of anterior Endo against EVL or embryo (top) and prechordal plate (bottom) against EVL or embryo. Per indicates percentage. As EVL cells can be specified spontaneously and are Nodal-independent^74^, here the proportions of prechordal plate and anterior Endo were adjusted by using the proportions of EVL cells as an internal control. (J) Venn plot showing the selection of Nodal direct target genes used for defining Nodal score (**see Methods**). (K) Dot plot visualizing Nodal score in wild-type, *ndr1*-morphant and *lft1*-mutant Nodal explants. Dot size indicates the percentage of cells within the cell group; the color indicates the average Nodal score levels. Each immunostaining experiment was performed for at least 3 independent replicates (technical replicates); more than 10 embryos were analyzed. Scale bar: 50 µm (E).

The results presented above suggest that differences in the Nodal-Lefty signaling may exist between PP and anterior Endo cells. It is widely accepted that Nodal-Lefty functions as reaction-diffusion system to facilitate Nodal gradient formation and cell fate determination^32–36^ (Figure 3D). As zebrafish possess redundant Nodal (Ndr1, Ndr2 and Ndr3) and Lefty (Lefty1 and Lefty2) paralogs, we firstly investigated the perturbation effects of individual Nodal or Lefty paralog on Nodal activity and its gradient. We generated an ectopic Nodal gradient in zebrafish animal pole cells by injection of *ndr2* mRNA at 128-cell stage in wild-type embryos, *ndr1* morphants and *lft1* mutants (Figures 3E, S6A-S6F and S6H-S6M). By immunostaining and quantification of the pSmad2 signaling, we observed a significant enhancement of Nodal activity in *lft1* mutants, while its level decreased upon *ndr1* knockdown (Figures 3E, 3F and S6A-S6M).

To further determine how Nodal-Lefty is involved in the regulation of PP and anterior Endo separation, we performed single-cell transcriptomic sequencing for Nodal explants constructed from *ndr1* morphants and *lft1* mutants at shield stage (Figure 3G). We integrated these two single-cell RNA-seq datasets with the dataset of wild-type Nodal explants^22^. After applying unsupervised clustering, we identified a total of 9 different cell clusters (Figures 3H, S7A and S7B). Among these clusters, two mesodermal clusters named mesoderm_PP and mesoderm_NT (Figure 3H) were obtained. Mesoderm_PP cells transcriptionally resembled the progenitors of PP evidenced by the high expression of *gsc* and *chrd*, while mesoderm_NT cells were the progenitors of notochord characterized by the high expression of *noto* and *tbxta* (Figures S7A and S7B). We observed that a small cluster of *sox32* and *sox17* positive anterior Endo cells was close to the PP cells on the UMAP (Figure 3H). We then calculated the proportions of PP and anterior Endo cells in the three datasets, and found that the proportions of both cell types increased in *lft1*-mutant explants, while decreased in *ndr1*-morphant explants compared to wild-type explants (Figures 3I and S7C).

To better estimate Nodal activity during PP and anterior Endo cell fate segregation, we defined a Nodal score by regressing out the expression level of all the Nodal downstream targets, which were differentially expressed upon Nodal treatment^22^ and also possessed pSmad2 binding peaks near their genetic loci^22,37^ (**see Methods**). A total of 29 Nodal downstream targets, whose expression exhibited a high correlation with Nodal activity, were used to calculate Nodal score (Figure 3J and S7L). We found Nodal score was dramatically decreased in *ndr1* knockdown samples and slightly increased in *lft1* mutant samples compared to that in wild-type samples (Figures 3K and S7D), which is consistent with the results of pSmad2 immunostaining (Figures 3E and 3F). We then examined the levels of Nodal score across all cell types in Nodal explants at 6 hpf, and found that axial mesendodermal cells, including PP, notochord and anterior Endo, displayed the highest Nodal scores (Figure S7E), which was consistent with previous knowledge that Nodal activity was the highest in the dorsal margin of zebrafish embryos at the onset of gastrulation^1,38,39^. A further comparison of Nodel scores between PP and anterior Endo revealed that PP had a sightly higher Nodal score than anterior Endo (Figure S7E). These analyses underscored the importance of Nodal signaling levels in PP and anterior Endo cell fate specification.

### A meticulous single-cell trajectory tree reveals the role of chromatin remodeling in PP and anterior Endo segregation

To thoroughly investigate the dynamic transcriptome changes during PP and anterior Endo separation, we reconstructed a single-cell trajectory tree using all PP and anterior Endo cells from three Nodal explant datasets at 6 hpf which was a crucial time point previously identified for the separation of these two cell trajectories (Figure 4A). This trajectory tree captured a continuous cell-state-transition process from PP progenitors to anterior Endo, and a key branching point in this process was identified (Figures 4A and S7F). To elucidate the molecular dynamics in this branching process, we performed differential expression analysis for these two cell trajectories around the branching point (Figure 4B). Three clusters of genes were identified, which were highly expressed in pre-branch (progenitors), left-branch (anterior Endo) and right-branch (PP) respectively. GO enrichment analysis using these three gene sets identified terms related to mesendodermal cell fate specification (Figure 4B). Interestingly, “chromatin organization” was significantly enriched in the pre-branch cells, and maintained high enrichment during PP specification, but was down-regulated upon endodermal fate specification (Figure 4B). Furthermore, the GO term “chromatin organization” was also enriched in the differentially expressed genes of the PP cells when comparing *lft1* mutant or *ndr1* knockdown samples to wild-type samples (Figures 4C-4G and S7G).

**Figure 4.**
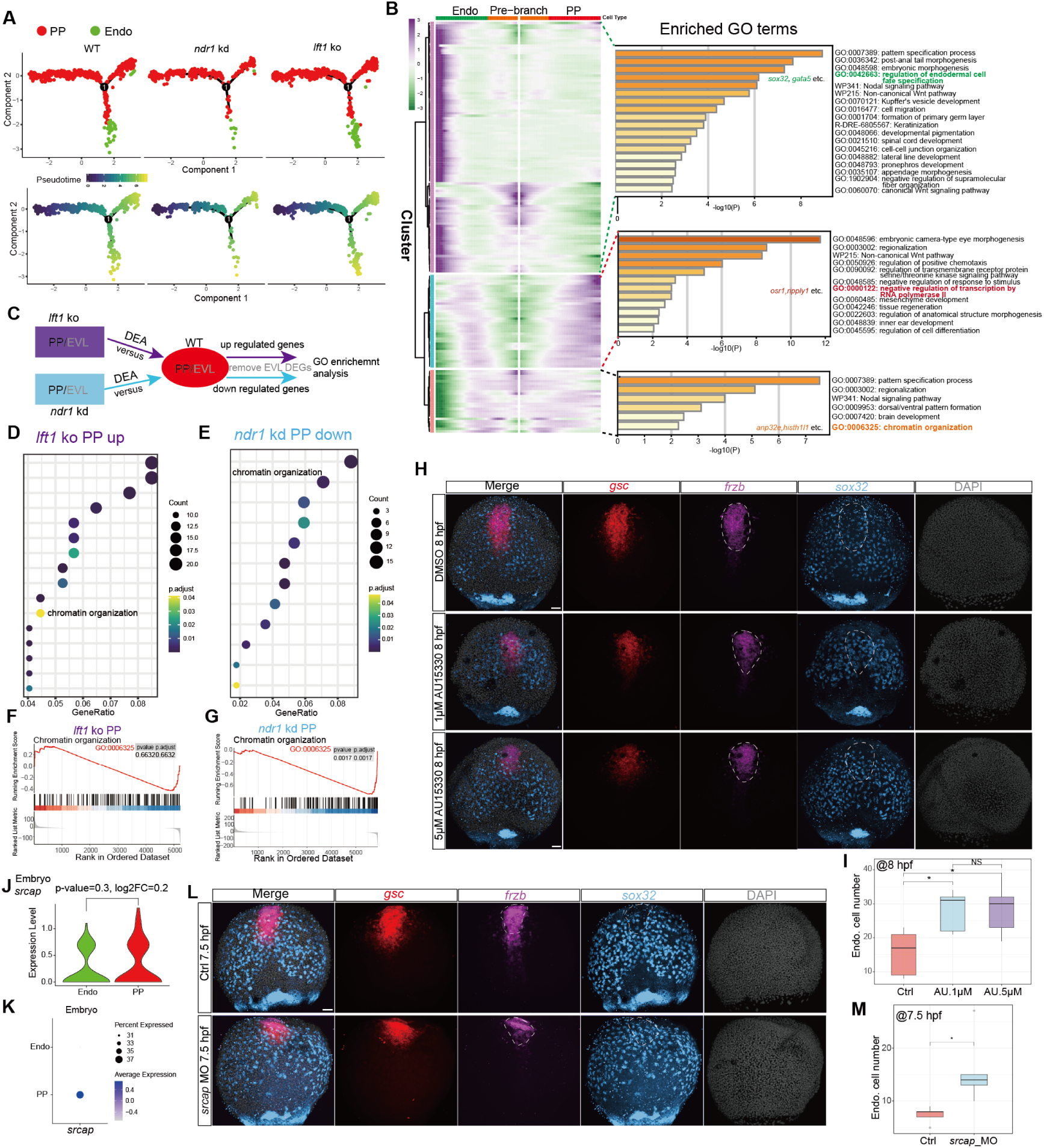
Single-cell trajectory tree analyses of PP and anterior Endo reveal that chromatin organization is involved in the separation of these two cell trajectories. (A) Pseudotime trajectory analysis of PP and anterior Endo using integrated single-cell datasets of wild-type (left), *ndr1*-morphant (middle) and *lft1*-mutant (right) Nodal explants. Cells are colored by cell types (top) and pseudotime levels (bottom). (B) Heatmaps representing clusters of genes that co-vary along the pseudotime during the separation of PP and anterior Endo from common progenitors. Top GO terms enriched by the genes in each cluster are listed with their corresponding adjusted P-values. Representative GO terms of each cluster are highlighted in different colors. (C) Diagram of gene set selection for GO analysis. As a control of batch effects, EVL differentially expressed (DE) genes are removed from mesoderm PP DE genes. (D and E) GO enrichment analysis of mesoderm PP up-regulated genes in *lft1* mutants (D) and down-regulated genes in *ndr1* morphants (E). Redundancy of enriched GO terms are removed, and each GO term contains at least 10 transcripts. Note that only the top 16 enriched GO terms identified in *lft1* mutant are plotted for a visualization purpose. (F and G) GSEA of “chromatin organization” term on PP cells from *lft1* mutants (F) and *ndr1* morphants (G) respectively. P-adjust indicates Benjamini-Hochberg adjusted p-value. (H-M) SWI/SNF complexes involved in regulating cell fate separation between the PP and anterior Endo. (H) HCR co-staining of *gsc*, *frzb* and *sox32* in wild-type embryos (top, n = 5/5, n: embryos were imaged, expression observed/total imaged), as well as those treated with 1 µM (middle, n = 4/5, n: embryos were imaged, expression observed/total imaged) and 5 µM (bottom, n = 5/6, expression observed/total imaged) AU15330. Regions framed by dotted white lines were used to quantify the cell number of anterior Endo. (I) Box plot showing the cell numbers of anterior Endo in (H). (J and K) Volin plot (J) and dot plot (K) indicating the expression levels of *srcap* in PP and anterior Endo cells in zebrafish embryos. (L) HCR co-staining of *gsc*, *frzb* and *sox32* in wild-type embryos and embryos with *srcap* knockdown. Embryos at 7.5 hpf (Ctrl, n = 6/6; *srcap* MO, n = 4/5; n: embryos were imaged, expression observed/total imaged) were evaluated. (M) Box plot showing the cell numbers of anterior Endo in (L). Statistical differences between two samples were evaluated by Student’s t-test (I and M). * indicates P-value < 0.05; NS indicates P-value >=0.05. Each HCR experiment was performed for at least 3 independent replicates (technical replicates); more than 40 (H) and 30 (L) embryos were analyzed. Scale bar: 50 µm (H and L).

These analyses unveiled a potential role of chromatin regulators in the regulation of PP and anterior Endo segregation, prompting us to identify the specific factors involved in this process. Among the candidates within GO term of “chromatin organization” (Figure 4B-4E), we identified a key member of SWI/SNF complex, *srcap*, whose expression is slightly higher in the PP than in the anterior endoderm cells in both embryos and Nodal explants (Figures 4J, 4K and S7H). Previous studies have demonstrated that the SWI/SNF nucleosome remodeling complex is necessary for TGF-β-induced transcription of numerous target genes^40,41^. To investigate the role of the SWI/SNF complex in the separation of PP and anterior Endo cell fates, we treated embryos with AU15330 (Figure 4H), a known degrader of SWI/SNF ATPase components^42^. Subsequently, we assessed the cell fate specification of PP and anterior Endo by HCR co-staining for *gsc*, *frzb* and *sox32*. We found that AU15330 treatment significantly increased the number of anterior Endo cells while decreasing the number of PP cells compared to DMSO-treated control embryos (Figures 4H, 4I and S7J). Similar effects were observed in *srcap* morphants, where the cell number of anterior Endo increased compared to wild-type embryos (Figures 4L, 4M, S7I and S7K).

Collectively, these findings underscore the importance of SWI/SNF complexes in ensuring accurate cell fate segregation within the anterior mesendoderm.

### Integrative analysis of single-nucleus RNA-Seq and ATAC-Seq reveals distinct chromatin states underlying differential expression profiling in PP and Endo cells

To delve deeper into the involvement of chromatin states in the separation of PP and Endo, we conducted single-cell assays for transposase-accessible chromatin (ATAC) and nuclear RNA sequencing in zebrafish embryos at 6 hpf (Figure 5A). Recognizing that accurate cell clustering cannot solely rely on chromatin accessibility dataset, we integrated this dataset with RNA expression data to achieve precise cell type identification. The resulting cells were clustered into 10 different groups on the UMAP (Figures 5B, S8A, and S8B).

**Figure 5.**
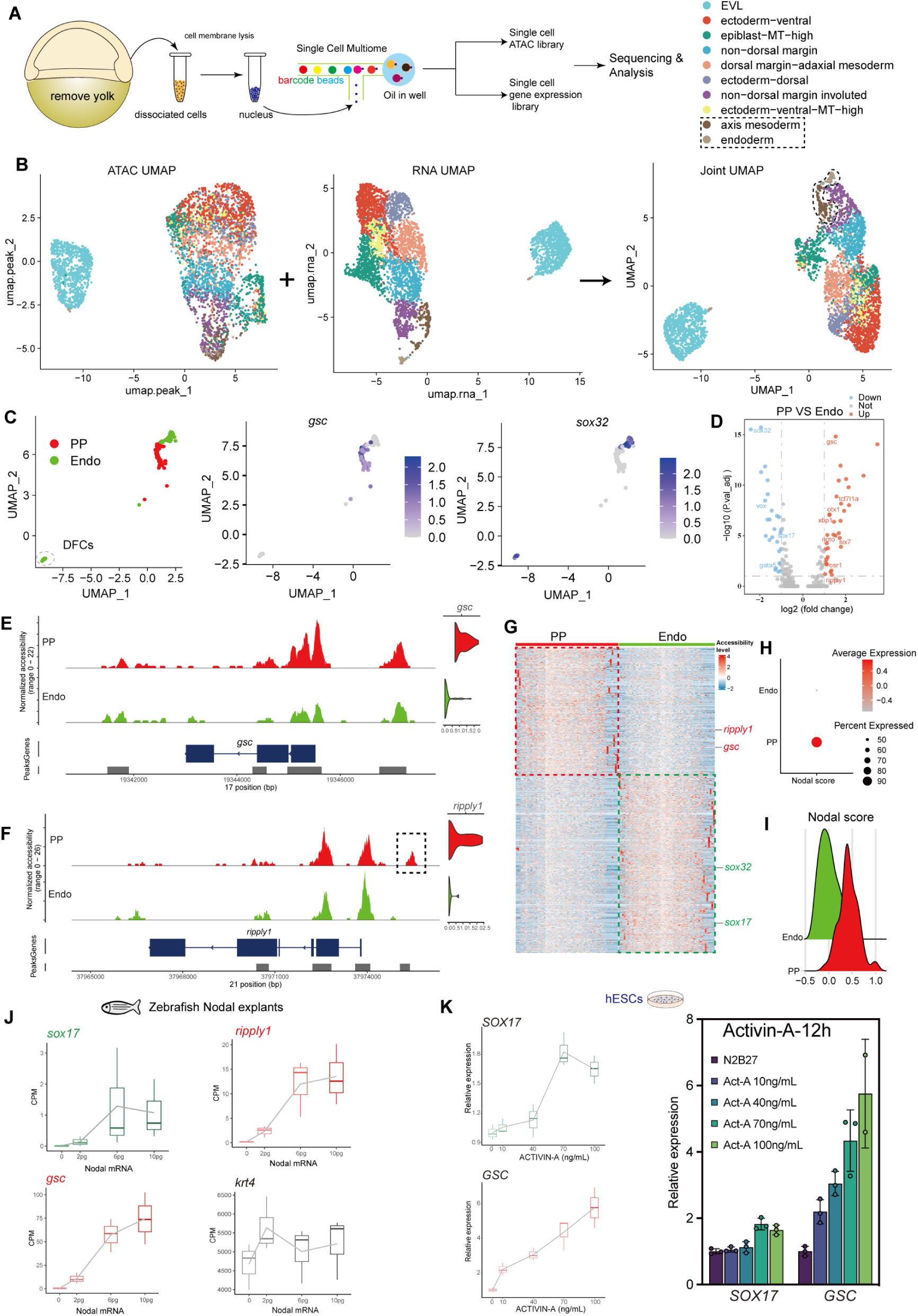
Investigating the mechanisms of PP and anterior Endo separation through integrative single-cell RNA-seq and ATAC-seq analysis. (A) Schematic diagram showing the experimental workflow of single-cell multi-omics of 6 hpf zebrafish embryos. (B) UMAPs based on scATAC-Seq (left), scRNA-Seq (middle) and co-embedding scRNA-seq and scATAC-seq (right) datasets. Cells are colored by different cell types. (C) UMAP plot of the PP and Endo cells. Cells are colored by cell types (left) and the expression of *gsc* (middle) and *sox32* (right). (D) Volcano plot showing the differentially expressed genes in PP and Endo based on RNA levels of single-cell multi-omics datasets. (E-F) Track plot showing chromatin accessibility of each cell type on the gene locus of *gsc* (E) and *ripply1* (F). (G) Heatmap showing the chromatin accessibility level of genes with differential chromatin accessibility in PP and Endo. Red and green box indicates genes with higher chromatin accessibility levels in PP and Endo cells respectively. (H and I) Dot plot (H) and ridge plot (I) showing the level of Nodal score in PP and Endo. (J) Box plots showing the expression profiles of *sox17* (Endo), *gsc* (PP), *ripply1* (PP) and *krt4* (EVL) in our previous bulk RNA-seq datasets of Nodal explants. (K) Box plots (left) and bar plot (right) showing the expression levels of *SOX17* and *GSC* in the human embryonic stem cells (hESCs) treated with different dosages of Activin protein (**see Methods**). The values were calculated as the mean ± SEM (K).

Using the marker gene *gsc*, we identified PP progenitor cells within the axis mesoderm, and then Endo and PP progenitor cells were selected for downstream analysis (Figure 5C). A small cluster of cells expressing *sox32*, *sox17* and *foxj1a*, which were located distantly from PP and Endo, were classified as dorsal forerunner cells (DFCs) and excluded from further analysis (Figures 5C and S8C).

Our data unveiled differential chromatin accessibility between PP and Endo cells (Figures 5G, S8K, and S8L). Correlation analysis revealed a strong correspondence between chromatin accessibility and RNA expression levels for marker genes in these two cell types (Figure S8D). To identify master regulators of PP and Endo separation, we performed differential analysis for both chromatin accessibility and gene expression (Figures 5D and S8E). A total of 170 up-regulated genes in PP, including *gsc* and *ripply1*, and 261 up-regulated genes in Endo, such as *sox32* and *sox17*, were identified. Most of these genes also exhibited differential chromatin opening states in PP and Endo fates (Figures 5E-5G and S8F-S8I).

Previous study reported that Nodal signaling can boost chromatin accessibility both *in vivo* and *ex vivo*^43,44^. And we observed that PP displayed a higher Nodal level compared to Endo cells (Figures S2J and S7E). In this ATAC and RNA-seq dataset, we also observed a higher Nodal score in PP cells (Figures 5H and 5I). Thus, it is possible that the differences in Nodal activity may lead to differential chromatin accessibility states and gene expression level in PP and Endo.

To further determine how Nodal signaling levels may differentially influence the gene expression dynamics, we first explored the expression profiles of key markers of anterior mesendoderm in the bulk RNA-seq data of Nodal explants injected with different dosages of Nodal mRNA^22^. We observed that the highest Nodal dosages did not result in further elevation of the expression of the endodermal marker *sox17* (Figure 5J), while higher concentrations of Nodal promoted the expression of *gsc* and *ripply1* (Figure 5J). Importantly, these patterns of differential marker expression in response to Nodal/Activin concentration were not only observed in our Nodal explant model but also validated in a human embryonic stem cell system (Figure 5K). These findings suggest that Nodal signaling levels play a conserved role in determining the cell fate specification, which influences the differentiation towards either Endo or PP fates. To further validate this finding, we injected different dosages of Nodal mRNA into one zebrafish animal pole blastomere at the 128-cell stage, and performed HCR co-staining for *sox32* and *frzb* at 6 hpf (Figure S9A). By quantifying the cell numbers of Endo and the area sizes of PP cluster, we found that moderate concentration of Nodal levels promoted Endo cell specification (Figures S9B-S9E).

Based on our aforementioned findings, it becomes evident that subtle variation in Nodal signaling levels may regulate PP and Endo gene expression. This variation is likely regulated by Nodal-Lefty regulatory loops, and contributes to the establishment of distinct chromatin states and transcriptional response of key transcriptional regulators within these two cell fates.

### Gsc and Ripply1 collaborate to suppress the specification of anterior Endo in zebrafish

It has been reported that endoderm differentiation could be suppressed by upregulation of *gsc*^18^, while the role of *ripply1* in endoderm development remains unknown, though it has been characterized as a transcriptional repressor ^45,46^. Moreover, both Gsc and Ripply1 are capable of inducing a secondary axis formation when overexpressed in the ventral side of zebrafish embryos^19,22^. Combining these findings from literature with our observation of high expression of *ripply1* in PP of Nodal explants and zebrafish embryos at 6 hpf (Figures 5D and S8I), we hypothesized that Ripply1 may also play a role in suppressing Endo specification. To test this hypothesis, we generated mutant lines for *gsc* and *ripply1* in zebrafish (Figure 6B). In order to comprehensively analyze the regulatory role of Ripply1 and Gsc in the specification of cell fates between PP and anterior Endo cells, we conducted a self-cross of the Tg(*gsc*^+/-^;*ripply1*^+/-^) lines (Figure 6A). Subsequently, we collected the descendant embryos to perform HCR co-staining for *frzb* and *sox32,* along with genotyping (Figure 6A). From the descendant embryos, we obtained a total of 9 genotypes with different phenotypes displaying variable defects in PP and anterior Endo separation (Figures 6C and S10A). Interestingly, we observed an increase in the number of anterior Endo cells with the loss of wild-type alleles of *gsc* and *ripply1* (Figures 6D, S10B, and S10C). To further investigate the mechanisms of Ripply1 suppressing anterior Endo specification in zebrafish, we injected *ripply1* mRNA at 1-cell stage in zebrafish embryos and found that Endo cell number was dramatically decreased as shown by whole-mount *in situ* hybridization (WISH) of *sox32* (Figure 6F). Additionally, we constructed a plasmid that contained *ripply1* cDNA downstream of the *sox17* promoter. Injection of this plasmid at 1-cell stage also resulted in a decrease in Endo specification shown by EGFP expression (Figure 6E). Taken together, these results demonstrated that Ripply1 acted as an Endo repressor in zebrafish.

**Figure 6.**
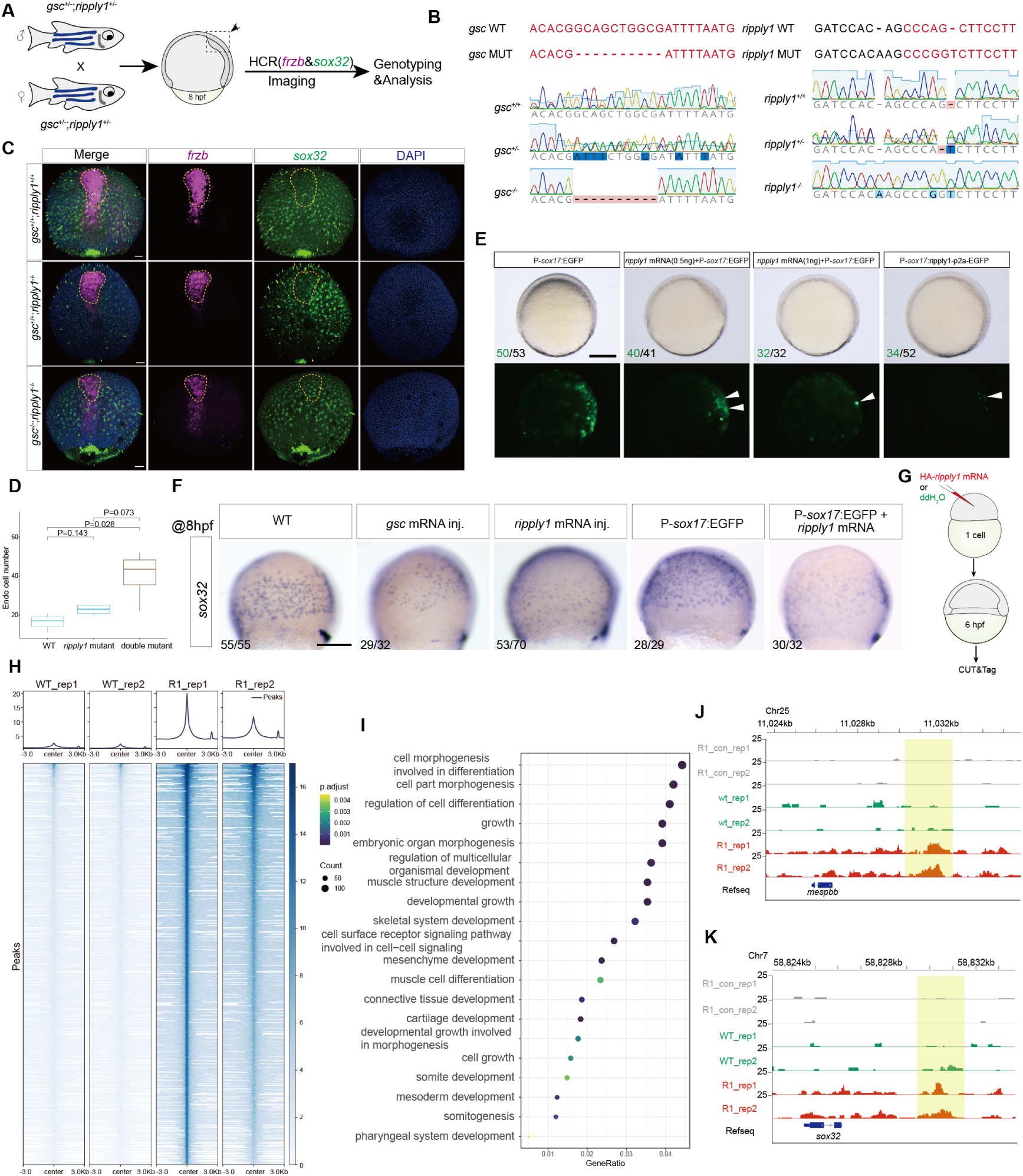
Exploring the mechanisms underlying the role of Ripply1 in the regulation of cell fate separation between PP and anterior Endo. (A) Schematic diagram illustrating the workflow for investigating the role of *gsc* and *ripply1* in mesendoderm cell fate separation through loss-of-function studies. (B) Schematic presentation of *gsc* and *ripply1* mutations and the sequencing results of each genotype. The mutations are a 10-bp deletion and a 1-bp insertion in *gsc* and *ripply1* coding sequences respectively. Texts in red indicate coding sequences. Sanger sequencing results display at the bottom, showcasing wild-type (top), heterozygous mutant (middle) and homozygous mutant (bottom) embryos of *gsc* and *ripply1* mutations. (C) HCR co-staining of *sox32* and *frzb* in the embryos of Tg(*gsc*^+/+^;*ripply1*^+/+^) (WT), Tg(*gsc*^+/+^;*ripply1*^-/-^) (*ripply1* mutant) and Tg(*gsc*^-/-^;*ripply1*^-/-^) (double mutant) at 8 hpf (Here embryos of three different genotypes are shown. Other genetypes are shown in Figure S10A.) [Tg(*gsc*^+/+^;*ripply1*^+/+^), n = 4/5; Tg(*gsc*^+/+^;*ripply1*^-/-^), n = 4/6; Tg(*gsc*^-/-^;*ripply1*^-/-^), n = 5/5; expression observed/total imaged]. (D) Box plot showing the quantification of the numbers of anterior Endo cells identified in (C). (E and F) Assessing the role of *ripply1* in anterior Endo specification. EGFP driven by *sox17* promotor (E) and WISH of *sox32* (F) are used to indicate the Endo cells. Embryos with *gsc* overexpression are used as a positive control. The statistics of observed/total sampled embryos were shown in bottom left of the panels (E and F). (G) Schematic diagram showing the strategy of conducting CUT&Tag experiment for *ripply1*. (H) Metaplots and heatmaps for CUT&Tag signals over R1_rep1 binding peaks (n=72,683) in wild-type and HA-*ripply1*-injected embryos. Two replicates were performed. (I) Dot plot showing the enriched GO terms using the genes that annotated from the differentially enriched peaks. (J and K) Screenshot of R1_con (technical control, without primary antibody, **see Methods**), WT (ddH_2_O-injected) and R1 (HA-*ripply1*-injected) CUT&Tag peaks around *mespbb* (J, positive control) and *sox32* (K) loci. Each HCR and ISH experiment was performed for at least 3 independent replicates (technical replicates). Statistical differences between two samples were evaluated by Student’s t-test (D). Scale bar: 50 µm (C) and 200 µm (E and F).

It is well established that Gsc functions as an Endo suppressor by directly binding to the promoter region of *sox17*^18^. However, the mechanism by which Ripply1 suppresses Endo specification remains unknown. To address this question, we injected HA-tagged *ripply1* mRNA into zebrafish embryos for conducting CUT&Tag experiments (Figures 6G, S11A and S11B). The CUT&Tag sequencing dataset was analysed following the standard process (Figures S11C-S11I), and 72,683 peaks were enriched in HA-*ripply1* injected samples (Figure 6H). Genes nearby the differentially enriched peaks were used to perform GO enrichment analysis (Figure 6I). Notably, we observed significant enrichment of terms related to “muscle structure development”, “skeletal system development” and “connective tissue development” (Figure 6I), which were consistent with the well-established functions of Ripply1 reported in the published studies^45–47^. Additionally, we also observed significant enrichment of GO terms such as “regulation of cell differentiation”, “negative regulation of signaling” and “negative regulation of transcription by RNA polymerase II” (Figure 6I), further supporting the transcriptional repressive function of Ripply1. More interestingly, Ripply1 binding peaks were enriched directly upstream of two key genes: *mespbb*, a known Ripply1 target, and *sox32*, a master regulator of endodermal specification (Figures 6J and 6K), suggesting a direct function of Ripply1 in the transcriptional regulation of the endodermal marker.

In summary, our data proposes a model within PP and anterior Endo cell fate separation. Variations of Nodal activities promote the establishment of distinct chromatin states in PP and anterior Endo fates, which is facilitated by SWI/SNF complexes. These differential chromatin accessibility patterns, in turn, drive the differential expression of key regulators such as *gsc* and *ripply1* in PP and anterior Endo. Collectively, these regulators collaborate to finely regulate the segregation of mesendoderm cell fates (Figure 7).

**Figure 7.**
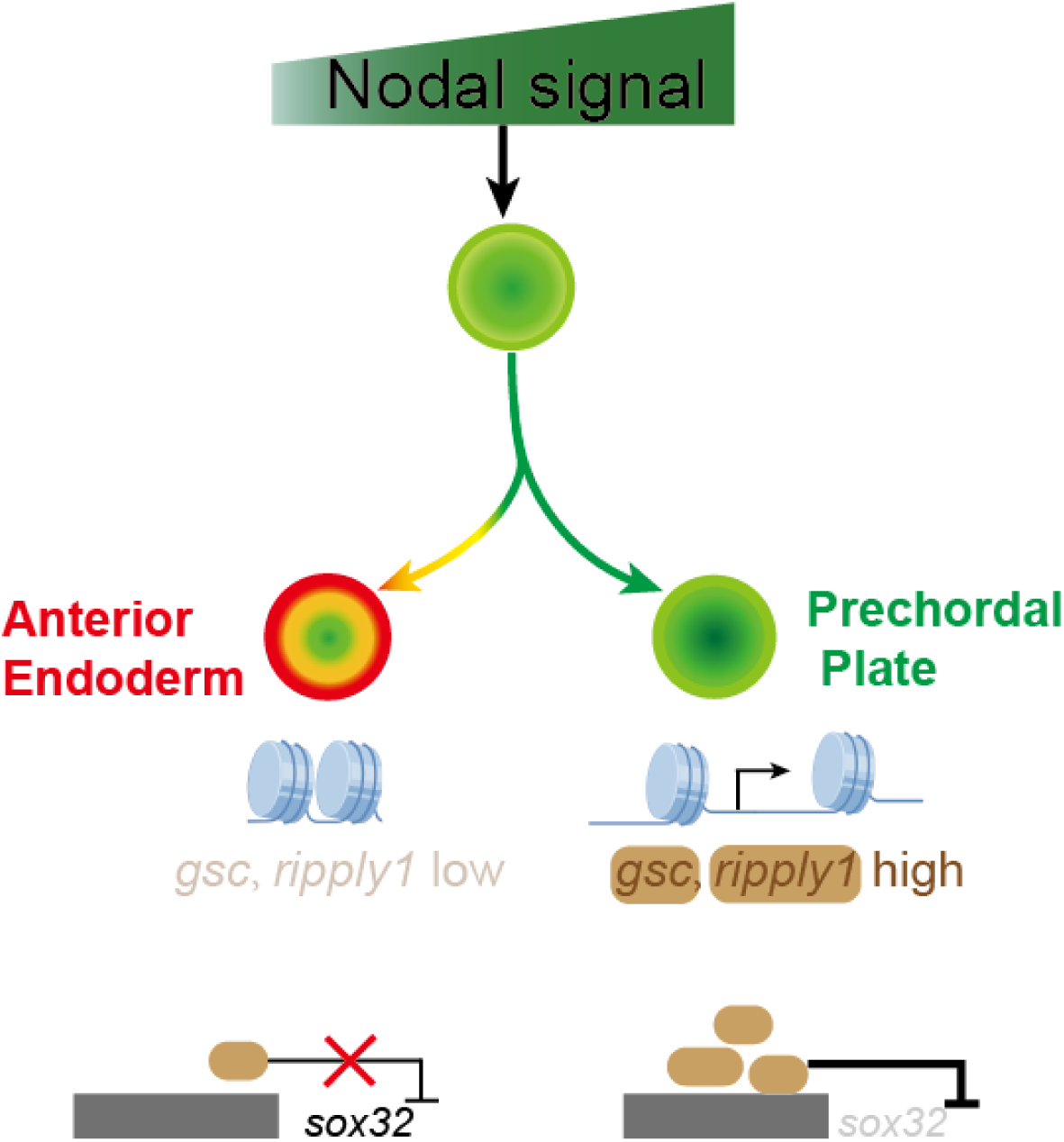
Model underlying the separation of PP and anterior Endo. An outline depicts PP and anterior Endo specification. Nodal signaling drives differential chromatin states between PP and anterior Endo through epigenetic regulators. And then, key transcriptional factors, such as *gsc* and *ripply1*, are differentially expressed in these two cell trajectories and suppress anterior Endo specification by directly binding to the *cis*-elements of *sox32*.

## Discussion

In this study, we delved into the intricacies of specifying and segregating PP and anterior Endo cell fates in zebrafish through single-cell multi-omics and live imaging analyses. Our findings revealed the origin of anterior Endo from PP progenitors and unveiled a correlation between subtle differences of Nodal signaling levels and fate commitment. These signaling variations were identified as potential regulators influencing the expression of cell-fate-specific genes at both epigenetic and transcriptional levels, with the collaboration of SWI/SNF complexes. Through gain-/loss-of-function studies, we demonstrated the collaborative suppression of Gsc and Ripply1 in anterior Endo specification.

While the induction of mesendodermal cell fates has been extensively studied^9,12,48,49^, the mechanisms underlying the separation of meso- and endo-fates from common progenitors remain a subject of ongoing research. The widely accepted notion posits that high Nodal signaling levels promote endoderm specification, while lower levels induce mesoderm^11–13^. These quantitative effects of Nodal signaling lead to differential activation of transcriptional targets which have differential sensitivity to Nodal activation^2,37^. However, recent studies challenge this perspective, proposing that Nodal signal level may not strictly determine cell fate separation of mesendoderm. Instead, Nodal signaling provides competency for mesendodermal progenitors to stochastically switch and commit to lineage-specific fates in zebrafish^16^. Moreover, the duration of Nodal signaling has been implicated in specifying different cell fates of mesendoderm, with prolonged signaling promoting prechordal plate specification and suppressing endoderm induction^18^. Besides, other important signaling pathways in early embryonic development also interact with Nodal signaling to regulate cell fate separation^13,31^. It has been well established that Fgf signaling plays a crucial role in regulating cell fate switching between endoderm and mesoderm in lateral margin of zebrafish embryo. In our current work, we preliminarily explored whether Fgf signaling plays roles in regulating cell fate segregation between PP and anterior Endo cells (Figure S12). We observed that several genes related to Fgf signaling were highly enriched in PP and anterior Endo cells (Figure S12B), and inhibition of Fgf signaling could obviously increase the cell number of anterior Endo (Figures S12A and S12C). The findings above suggest that Fgf signaling inhibits the specification of anterior Endo from PP progenitors in zebrafish, which is a similar process to the suppression observed during lateral endoderm specification. However, further investigation is required to understand the specific interactions between Fgf signaling^50^ and factors such as Gsc, Ripply1, Sox32^51,52^ and the SWI/SNF complexes, as well as their roles in regulating the fate diversification of PP and anterior Endo fates.

Our study contributed a new layer of understanding to the regulation of mesendodermal cell fate segregation by Nodal signaling. We proposed that Nodal signaling may influence chromatin remodeling factors, thereby affecting the chromatin accessibility states of key regulators such as *gsc* and *ripply1*. It is worth noting that *gsc* and *ripply1* have been identified as direct targets of Nodal signaling^22^. This suggests that the transcription of these genes may directly and sensitively respond to variations in Nodal signaling levels. Based on this understanding, we hypothesize that Nodal signaling can achieve more precise and robust control over the fate segregation of closely related cell types by introducing an additional layer of epigenetic regulatory mechanisms. Additionally, our findings underscored the pivotal role of chromatin openness in morphogen interpretation during development, emphasizing the need to consider chromatin structure when studying how morphogen gradients regulate patterning formation across different species.

In our work, combining single-cell multi-omics analyses with perturbation experiments, we revealed that chromatin remodeling together with Nodal morphogen promoted the PP and anterior Endo separation in zebrafish. However, which and how chromatin remodeling factors respond to Nodal signaling to facilitate fate diversification remains largely unknown. Our current study identified a key member of SWI/SNF complexes, *srcap*, whose expression was slightly higher in the PP than in the anterior endoderm cells. What triggers the transcriptional difference of *srcap* in these two cell types is still an open question. One of the hypotheses is that the transcription of *srcap* is very sensitive to Nodal signaling (Nodal concentration or Nodal duration), and a little variance in Nodal signaling can drive differential activation of *srcap*. Another speculation is that the activation of *srcap* needs the assistance of pioneer factors, like forkhead family^40,43^, whose binding motifs are particularly accessible in PP cells compared to anterior Endo cells (Figures S8K and S8L). All these hypotheses need to be further determined in future works.

Despite these insights, several questions remain unanswered. Firstly, we discovered a potential role of Nodal activity levels in the separation of PP and anterior Endo cells. However, further investigation is required to understand how Nodal activity levels and signaling duration are integrated to regulate this process. Secondly, although our study determined the collective role of Gsc and Ripply1 in suppressing endoderm specification, the potential involvement of other unidentified transcriptional repressors, as suggested in previous work^18^, warrants further investigation. Notably, even in double homozygous mutants of *gsc* and *ripply1*, the specification of PP clusters was still detected, and the enrichment of another transcriptional repressor, *osr1*, in PP cells (Figure S8J) suggested a complex network of collective endoderm repressors in preventing cells from differentiating into anterior Endo. Thus, it would be worth studying how Gsc, Ripply1, Osr1 and other potential factors collectively regulate PP and anterior Endo specification in the future. Lastly, the anterior endoderm and ventral-lateral endoderm were mixed together in our single-cell ATAC-seq dataset, which brought in interferences when we interpreted the mechanisms of the separation between PP and anterior Endo. More replicates of embryonic single-cell multi-omics or performing single-cell multi-omics on Nodal explants may help to address this issue in future studies.

## Methods

## Animal ethics

Zebrafish strains were conducted following standard procedures, and experimental procedures were approved by the Institutional Review Board of Zhejiang University. The protocol number is ZJU20220375.

### Generation of Nodal explants

Nodal explant was generated as our previous work described^53^. 10 pg *ndr2* mRNA was injected into one blastomere of zebrafish embryonic animal pole at the 128-cell stage. About half of the animal pole region of the embryo was cut off from the blastula at the 1k-cell stage, and was cultured in a Petri dish coated with 1.5% agarose gel filled with Dulbecco’s modified Eagle’s medium/nutrient mixture F-12 (DMEM/F-12) with 10 mM HEPES, 1x minimum essential medium (MEM) containing non-essential amino acids and supplemented with 7 mM CaCl_2_, 1 mM sodium pyruvate, 50 μg/ml gentamycin, 100 μM 2-mercaptoethanol, 1× antibiotic-antimycotic (15240062, Thermo Fisher) and 10% serum replacement. For generating *ndr1*-morphant Nodal explants, *ndr1* morpholino (0.5 mM, 2 nL) was injected into yolk of wild-type zebrafish embryos at 1-cell stage; while for generating *lft1*-mutant Nodal explants, the embryos of *lft1* mutants were used to construct Nodal explants. The sequence of *ndr1* morpholino was 5’ - ATGTCAAATCAAGGTAATAATCCAC - 3’.

### Live imaging

Tg(*gsc*:EGFP;*sox17*:dsRed) embryos and Nodal explants generated from those embryos were used to perform live imaging analysis. 0.5% low-melt agarose was used to mount the embryos or Nodal explants in glass-bottom dishes (Cellvis, D35-20-0-N). The embryos or explants were live-imaged by confocal laser scanning microscopy (OLYMPUS FV12-IXCOV) using a 20X objective lens or a 40X oil immersion lens.

### *In situ* hybridization chain reaction (HCR)

HCR co-staining of *frzb*, *sox32* and *gsc* was performed as previous studies described^21^. The probes of those genes were ordered and produced from Molecular Technologies (http://www.moleculartechnologies.org/). Embryos were fixed in DEPC-treated PBS with 4%(w/v) paraformaldehyde at 4°C overnight. The embryos were incubated in 500 μL 30% formamide probe hybridization with 2 pmol of each HCR probe set at 37°C overnight. Next day, 30% formamide probe wash buffer was used to stop hybridization by repeated washing of the embryos at 37°C. Fluorescent signals were generated and amplified by probes that bound to fluorescent HCR amplifiers in an amplification buffer overnight at room temperature. To halt this process, the samples were washed several times using 5× SSC with 0.001% Tween 20. The reaction buffers, fluorescent hairpins and probes were manufactured by Molecular Technologies. OLYMPUS FV12-IXCOV confocal microscope was used to photograph those HCR co-staining samples.

### Double-color RNA-fluorescence *in situ* hybridization (FISH)

Double-color FISH of *gsc* and *sox17* was conducted according to the protocol from manufactory (http://www.pinpoease.com/, GD Pinpoease Biotech Co., Ltd.). Wild-type embryos or Nodal explants (6 hpf, 7 hpf and 8 hpf) were fixed in DEPC-treated PBS with 4%(w/v) paraformaldehyde at 4°C overnight. Pre-A solution was used to inhibit the peroxidase activity. Target RNA molecules were exposed through protease treatment, followed by hybridization with probes at 4°C for 2 hours. The fluorescent signal was amplified through sequential reactions 1, 2, and 3. Lastly, a tyramide fluorescent substrate (OpalTM520, Akoya Biosciences) was used to incubate the embryos, enabling fluorescent labeling of the target RNA through the Tyramide Signal Amplification (TSA) assay.

### Immunostaining and quantification of pSmad2

Samples were fixed at 4% paraformaldehyde overnight for pSmad2/3 immunostaining. The experiment was performed as previously described^53^. Confocal laser scanning microscopy (OLYMPUS FV12-IXCOV) was used to scan the immunostained samples. Imaris (Version 9.7) software was used to quantify the signaling activity of pSmad2 (488 nm), RFP (561 nm) and DAPI (405 nm). Firstly, the raw image data was loaded in Imaris and visualized in a three-dimensional (3D) view. The Spot plugin from Imaris was used to detect spots with pSmad2 signal. The diameter of spot detection was set to 5-6 μm for GFP channel. The spots were automatically identified with default parameters. And then, as the identification of some spots was inaccurate, we manually added or removed some spots. Lastly, the coordinates and signal intensities of each spot were exported for next-step analyses. To make all images comparable, we used DAPI signal as an internal control to normalize the signal intensity in other channels. When plotting a pSmad2 signal intensity gradient along a distance, the cell with the highest pSmad2 signal was assigned as the start point, and then the distance of other cells was calculated by comparing them with the start point cell. The fitting curve was plotted by ggplot2^54^ in R (https://www.r-project.org).

### Whole-mount *in situ* hybridization (WISH)

The processes of WISH and the construction of *sox32* probes were described in our previous work^53^. Embryos were fixed in DEPC-treated PBS with 4%(w/v) paraformaldehyde at 4°C overnight. To enable long-term storage, the embryos were dehydrated using 100% methanol. For WISH experiments, probes were labeled with digoxigenin-11-UTP (Roche Diagnostics), and the substrate was NBT/BCIP.

### Activation of Activin/Nodal signaling in human embryonic stem cell system

Human embryonic stem cell (hESC) lines were used between passage 45-50. Cells were seeded equally on matrigel-coated plate and cultured in mTeSR medium. Before the experiment, each group was pretreated in N2B27 medium for 6 hours. Then the control group was maintained in N2B27, while the experiment groups were cultured in N2B27 medium supplemented with different concentrations of Activn-A (0, 10, 40, 70 and 100 ng/mL). The cellular RNA of each group was extracted with Trizol after 12 hours of treatment.

### Overexpression of *gsc* and *ripply1* in zebrafish embryos

The mRNA of *gsc* and *ripply1* (synthesized *in vitro*) and *sox17* promotor-driven *ripply1* plasmid were co-injected into zebrafish embryos at 1-cell stage. The ddH_2_O was injected as control. The embryos were imaged or fixed at 8 hpf. The *sox32* probe was used for WISH.

### Generating heterozygous embryos of *gsc* and *ripply1*

The *ripply1* mutants were obtained from Dr. Ming Shao’s lab at Shandong University. We knockout *gsc* in *ripply1*-mutant embryos by CRISPR-Cas9 system. F0 embryos were raised up, and then crossed with wild-type (AB). The descendant embryos (F1) were raised up and genotyped. The adult zebrafish of the target genotype were selected.

### Quantifying the cell number of anterior Endo

Imaris (version 9.7) was used to identify the anterior Endo cells based on the HCR images. Briefly, the raw image data was imported into Imaris, and the object was visualized in a 3D view. The Spot plugin from Imaris was used to detect each cell. The diameter of spot detection was set to 10-12 μm for GFP channel (staining for *sox32* or *sox17*). The candidate spots were selected by setting an appropriate quality threshold. Finally, the coordinates and signal intensities of each spot were exported from Imaris and used for further analysis.

### CUT&Tag sample preparation

HA-*ripply1* sequence with homologous sequences was generated through two PCR steps using two separate sets of primers (Table S1, reverse primer was common, *ripply1*-F1 primer: tcaaggcctctcgagcctctagaATGTACCCATACGATGTTCCAGATTACGCT; *ripply1*-F2 primer: ACGATGTTCCAGATTACGCTGGCAGCATGAATTCTGTGTGCTTTGCCACT; *ripply1*-R primer: TACGACTCACTATAGTTCTAGAtcagttgaaagctgtgaagtga). The details of generating mRNA *in vitro* was described in our previous protocol^21^. Briefly, the HA-*ripply1* PCR products were inserted into pCS^2+^ plasmid through homologous recombination using <u>ClonExpress II One</u> <u>Step Cloning Kit</u>. The plasmid containing HA-*ripply1* was linearized using *Not1* cleavage, and HA-*ripply1* mRNA were generated through mRNA *in vitro* synthesis using <u>mMESSAGE</u> <u>mMACHINETM SP6 transcription kit</u>. Wild-type embryos were injected with 500 pg HA-*ripply1* mRNA or ddH_2_O at one-cell stage and harvested at 6 hpf. Yolk was removed through pipetting, and the cells were collected. The cell viability was assessed by Trypan blue staining and the cell number was then counted.

### CUT&Tag experiment

CUT&Tag was performed on samples of wildtype, HA-*ripply1*-injected and HA-ripply1-injected without primary antibody using <u>Hyperactive Universal CUT&Tag Assay Kit for</u> <u>Illumina Pro</u> (Vazyme, TD904), and each sample had two replicates. Briefly, around 100,000 cells were collected and immobilized on Concanavalin A-coat Magnetic Beads. Cells were then incubated with primary antibody (HA-Tag Rabbit mAb, Cell Signaling Technology, 3724S, 1:50 dilution) at 4°C for 12 h-16 h. Tethered cells were washed thoroughly to remove excessive primary antibody and incubated with secondary antibody (Goat Anti-Rabbit IgG H&L, Vazyme, Ab207-01, 1:100 dilution) for 60 minutes at room temperature. Then samples were washed thoroughly to remove unbound secondary antibody and incubated with pA/G-Tnp for 1 hour at room temperature. DNA was then fragmentated through Trueprep Tagment Buffer L (TTBL). Fragmented DNA was extracted using DNA extract beads and amplified through PCR. PCR products were purified using VAHTS DNA CLEAN Beads (Vazyme, N411) and then sent to Novogene Co., Ltd. (Beijing, China) for sequencing. Paired-end sequencing at 150 bp read length was performed on an NovaSeq 6000 System.

### CUT&Tag analysis

Sequencing data of CUT&Tag samples were first treated by Cutadapt v4.5^55^ to remove adapters and trim reads using the following parameters “*-m 18 -q 30,30 --max-n=0.05 -e 0.2 -n 2*”. The trimmed reads were then aligned to zebrafish genome (danRer11) through Bowtie2 v2.5.2^56^ using the following parameters “*--end-to-end --very-sensitive --no-mixed --no-discordant --phred33 -I 10 -X 700*”. Samtools v1.18^57^ and BEDtools v2.31.0^58^ were then used for post-alignment processing to covert file formats in preparation for the peak calling and visualization. BigWig tracks were generated for visualization of chromatin landscapes in interested regions using the Integrative Genomics Viewer (IGV)^59^. To assess the reproducibility between replicates, the genome was spilt into 500 bp bins, and a Pearson correlation of the log_2_-transformed read counts in each bin was calculated between datasets.

SEACR^60^ was used to call “non stringent” peaks for wild-type and HA-*ripply1*-injected samples using merged HA-*ripply1*-injected samples without primary antibody as control. Additionally, top 1% of enrich regions were selected as peaks by AUC without controls. Peaks were annotated by ChIPpeakAnno v3.34.1^61^ using UCSC annotation from TxDb.Drerio.UCSC.danRer11.refGene v3.4.6^62^. Significantly enriched peaks in HA-*ripply1*-injected samples were identified using DESeq2 v1.40.2^63^, and enriched GO terms were identified using clusterProfiler v4.9.0.2^64^. Signal intensity heatmaps were generated using deeptools v3.5.4^65^. The peaks of HA-*ripply1*-injected rep1 were used as the reference to generate computeMatrix using the *reference-point* model (*--skipZeros --afterRegionStartLength 3000 --beforeRegionStartLength 3000 --referencePoint center*), and *PlotHeatmap* function was used to plot signal intensity. To obtain more convincible Ripply1-binding targets, we annotated all peaks enriched in the treatment group and performed differential analysis, selecting genes with log₂FoldChange > 3, padj < 0.5, and baseMean > 30 as candidate targets of Ripply1 binding.

### Processing and analyzing single-cell RNA-seq data

Illumina sequencing reads were aligned to the reference genome (GRCz11) of zebrafish through 10× Genomics CellRanger pipeline (version 3.0.2) with default parameters. Nodal explants generated from *lft1* mutants and *ndr1* morphants yielded 5,419 and 5,195 cells respectively. The expression matrix of each sample was obtained via CellRanger pipeline analysis and used for further analysis. The single-cell RNA-seq data of wild-type Nodal explants was reused from our previous study^53^.

### Processing and analyzing single-cell multi-omics data

Illumina sequencing reads were aligned to the reference genome (GRCz11) of zebrafish using 10× Genomics CellRanger-arc pipeline (version 2.0.0) with default parameters. Expression matrix and fragments files were obtained after running the pipeline. 4,879 cells were obtained. Signac^66^ was used for downstream analyses with default parameters. Low-quality cells were filtered out with nCount_ATAC < 100000 & nCount_RNA < 50000 & nCount_ATAC > 1000 & nCount_RNA > 1000 & nucleosome_signal < 2 & TSS.enrichment > 1 & percent.mt < 4. And then, gene expression data and DNA accessibility data were processed respectively by Signac with default parameters. The RNA expression level was utilized to identify cluster markers for cell annotation. Gene activity matrix was obtained by assessing the chromatin accessibility associated with each gene through *GeneActivity* function.

### Differential expression analysis of PP cells between wild-type, *lft1*-mutant and *ndr1*-morphant Nodal explants

The three sing-cell RNA-seq datasets of Nodal explants constructed from wild-type embryos, *lft1* mutants and *ndr1* morphants were integrated by Seurat^67^. To investigate the functional and regulatory effects of Nodal signals on cell differentiation, we compared the wild-type mesoderm PP cells to the mesoderm PP cells in *lft1* mutants and *ndr1* morphants respectively, and differentially-expressed (DE) genes were identified through Seurat v4.0.2 (https://github.com/satijalab/seurat) with a Bonferroni adjusted p-value < 0.05. As it is known that EVL is not affected by Nodal signaling^53,68^, DE genes identified in EVL were regarded as background noises and were removed from the mesoderm PP DE genes. The mesoderm PP DE genes positively correlated with Nodal concentration were then selected, constructing two gene sets: upregulated DE genes in *lft1* mutants and downregulated DE genes in *ndr1* morphants. GO enrichment analysis was performed on these two gene sets respectively using clusterProfilter v4.2.2 (https://github.com/YuLab-SMU/clusterProfiler). The enriched GO terms in “biological process” subontology were identified using a Benjamini-Hochberg adjusted p-value < 0.05.

### Cell-cell communication analysis by LIANA

R package, LIANA^30^ v0.1.5 (https://github.com/saezlab/liana/), was used for the investigation of ligand-receptor interactions among different cell types. The ligand-receptor analysis was performed against the ligand-receptor reference achieved from Omnipath resource using 5 different methods implemented in LIANA, including Natmi, Connectome, SingleCellSingalR, iTALK and CellPhoneDBv2. The consensus rank aggregated from the results of these methods was used for the identification of enriched ligand-receptor interactions.

### Inferring cell-cell communication by CellChat

CellChat^69^ was employed to systematically analyze cell-cell communication based on prior known zebrafish ligand-receptor interaction database CellChatDB. As some known interactions were missing in the database, we manually added some interactions of interest into the database. The expression data of scRNA-seq was preprocessed to identify over-expressed ligands and receptors for each group. CellChat inferred communication probability between two interacting cell groups based on the average gene expression of a ligand in one cell group and the average gene expression of a receptor in another cell group. The communication probabilities of signaling pathways were then calculated by summarizing the communication probabilities of all ligands-receptors interactions associated with each pathway.

### Gene set enrichment analysis

The wild-type mesoderm PP gene expression was compared to the mesoderm PP gene expression of *lft1* mutants and *ndr1* morphants respectively using Seurat v4.0.2 (https://github.com/satijalab/seurat). The genes expressed in more than 10% cells in either of the two compared populations were retained. A ranked list was formed on the retained genes using sign(log_2_FC) * (-log_10_PValue) as the ranking statistic. The mapping between GO terms and zebrafish genes was achieved through org.Dr.eg.db v3.14.0 (https://bioconductor.org/packages/org.Dr.eg.db/). GSEA was performed on the selected GO terms by clusterProfiler v4.2.2 (https://github.com/YuLab-SMU/clusterProfiler), using fgsea (https://github.com/ctlab/fgsea) with 10^5^ interactions. R package, enrichplot v1.14.2 (https://github.com/YuLab-SMU/enrichplot), was then used to visualize the GSEA results.

### Single-cell RNA-seq trajectory analysis

Three different methods were used to perform single-cell RNA-seq trajectory analysis. The developmental trajectory tree of zebrafish embryos and Nodal explants was constructed by URD^70^ with default parameters as described in previous studies^70^. The branchpoint preference plot of PP and anterior Endo was constructed by URD with default parameters. Firstly, *branchpointPreferenceLayout* function was used to define the preference layout for the branchpoint. And then stage, pseudotime information and gene expression were plotted on the branchpoint using *plotBranchpoint* function. The expression matrix of PP and anterior Endo cells was also used to construct trajectory tree by monocle2^27^ and monocle3^71^ with default parameters. And differential expression analysis was performed at the branching point of PP and anterior Endo separation on trajectory tree constructed by *BEAM* function in monocle2. Modules of co-regulated genes along PP or anterior Endo differentiation trajectory were identified by *find_gene_modules* function in monocle3 with default parameters. The cells contributed to the preference layout for the Endo and PP brachpoint from URD analysis were also selected as inputs for Palantir^26^. We used Harmony^72^ to calculate the augmented affinity matrix for data of all time points. Data was visualized using force directed layouts. Palantir was used for downstream analysis, and a *nanog^high^*cell was used as the start cell. A *frzb^high^* cell (PP) and a *sox17^high^* cell (Endo) were selected as the terminal cells. Anterior Endo and PP cell differential trajectory was detected by *palantir.core.run_palantir* function from Palantir. And finally, gene expression trend along Palantir inferred pseudotime was calculated by *palantir.presults.compute_gene_trends* function.

### Calculating Nodal score in single-cell RNA-seq data

Nodal functions through activating a set of key downstream genes in a concentration-dependent manner. Our previous study identified 105 Nodal immediate target genes by analyzing bulk RNA-seq datasets of Nodal mRNA injected explants^22^. We overlapped these 105 genes with 61 Nodal direct targets, which were previously identified through ChIP-seq analysis of pSmad2^37^. This analysis allowed us to pinpoint 29 Nodal downstream genes whose expression was highly sensitive to Nodal activity. To quantify Nodal activity, we defined a Nodal score based on its downstream transcriptional response. This score was calculated by regressing out the expression levels of these 29 Nodal downstream genes. To accomplish this, the *AddModuleScore* function in Seurat was employed.

## Supporting information

DataS1

DataS2

DataS3

DataS4

DataS5

DataS6

DataS7

DataS8

## Data availability

The data generated in this study is deposited in the Gene Expression Omnibus (GEO). The single-cell RNA sequencing and single-cell multi-omics data is under accession number GSE223636, and the CUT&Tag data is under the accession number GSE249292.

## Code availability

Any custom code and data are available from the authors upon request. All analyses are based on previously published code and software.

## Additional information

Supplemental Information: Supplementary Figures 1-12, Supplementary Table S1, Supplementary Data 1-8 and Supplementary Movies 1-5.

## Acknowledgments

We would like to thank Dr. Alex Schier for his kind gift of the *lft1* mutant. We thank Dr. Ming Shao from Shandong University for his kind gift of the *ripply1* mutant. We thank Dr. Peng Xia from Zhejiang University for his kind gift of the Tg(*gsc*:EGFP) embryos. We also thank Dr. Nai-He Jing at University of Chinese Academy of Sciences, Dr. Jun Ma, Dr. Xiao-hang Yang, Dr. Min-Xin Guan, Dr. Feng He and the members of Laboratory of Development and Organogenesis (LDO) at Zhejiang University for helpful suggestions and discussions. We thank Mr. Guang-Xu Zhang from Biostar Technology for the technical help on constructing 10x single-cell multi-omics sequencing library. We thank Shuang-Shuang Liu from the Imaging Platform and Ying-Niang Li from the zebrafish core facility at Zhejiang University School of Medicine for their technical support. This work was supported by grants from the Chinese National Key Research and Development Project (2022YFA1103100, 2019YFA0802402), the National Scientific Foundation of China (32050109, 32300688, 32300677) and the China Postdoctoral Science Foundation (2024M752767).

## Author Contributions

PFX, TC and HQL conceived and designed the research; TC, XL, YD, YMT, YYX, CYC, CL, YFL, YH, DHZ, XX, XXF, ZXJ, JYL and JM performed research; TC and YD analyzed the single-cell RNA-seq and single-cell multi-omics data; YD performed the CUT&Tag experiment and analyzed the data; TC, YD, HQL and PFX wrote the manuscript; all authors reviewed and approved the manuscript. PFX and HQL supervised this study. Fundings for the study were provided by PFX, TC and YD.

