## Supplementary Information for "Ripply1 and Gsc collectively suppress anterior endoderm differentiation from prechordal plate progenitors"

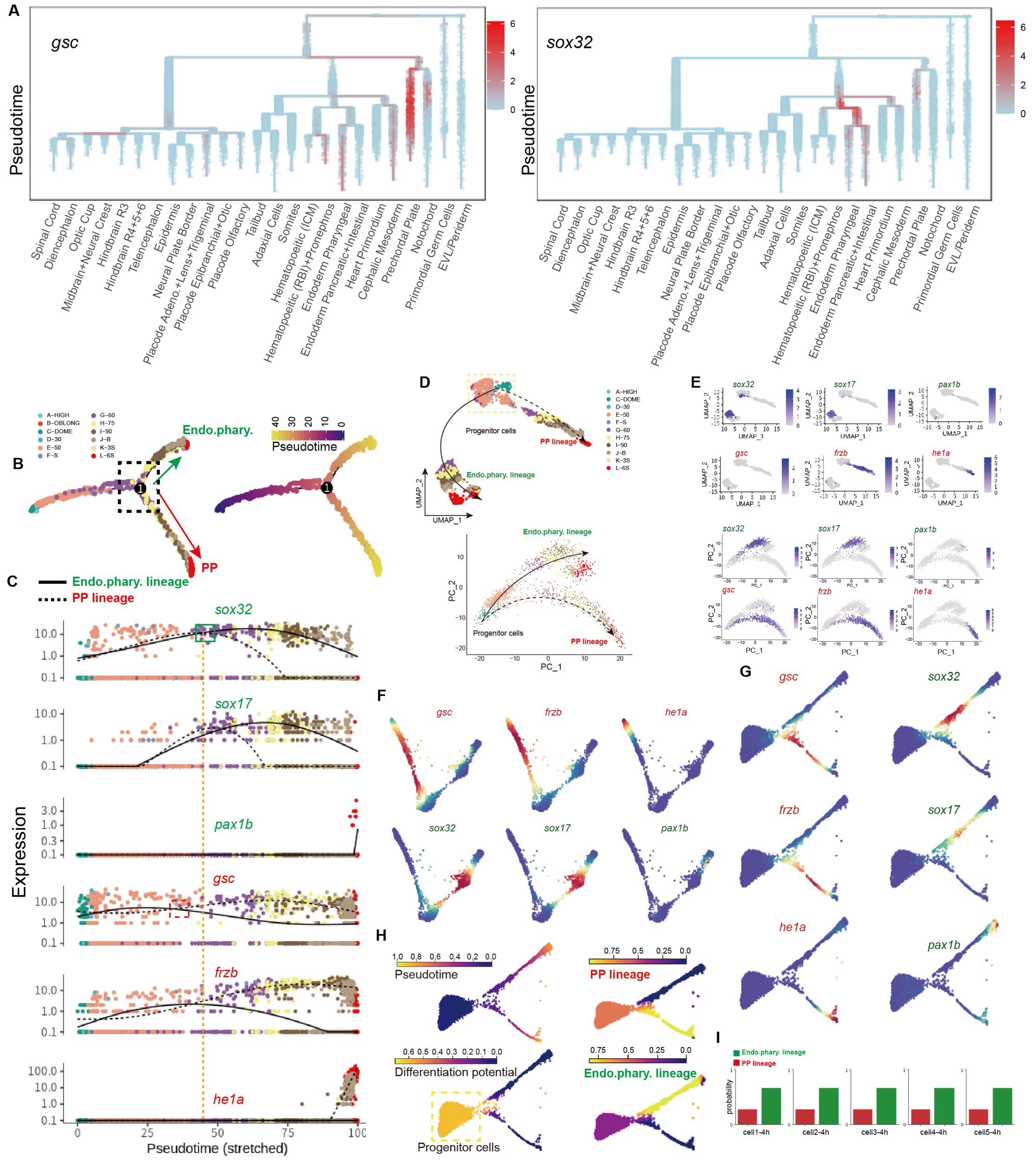


**Figure S1 Investigating the mechanisms of prechordal plate and anterior endoderm separation by zebrafish embryonic single-cell datasets.** (A) URD plot showing the expression of *gsc* and *sox32* on the zebrafish single-cell trajectory tree. Cells are colored by the expression level. (B) Single-cell pseudotime-ordered trajectory analysis of prechordal plate and anterior endoderm. The arrows indicate the possible differentiation direction. The black dotted frame indicates the branching point of these two cell trajectories. Cells are colored by developmental stages (left) and pseudotime (right). (C) Expression dynamics of prechordal plate and anterior Endo markers along pseudotime axis inferred by Monocle[1]. The green frame and the dotted red frame respectively indicate the points that *sox32* (also indicated by the orange dotted line) and *gsc* start to be differentially expressed in these two cell trajectories. (D) UMAP[2] (up) and PCA plot (bottom) showing the cell maps of prechordal plate and anterior endoderm during early development in zebrafish. Cells are colored by developmental stages; the solid and dotted arrows indicate the inferred differentiation trajectories of prechordal plate and anterior endoderm respectively; the yellow dotted frame shows the progenitor cells of these two cell trajectories. (E) The expression of *gsc*, *frzb*, *he1a*, *sox32*, *sox17* and *pax1b* on UMAP (up) and PCA plot (bottom) of prechordal plate and anterior endoderm. (F) The expression of *gsc*, *frzb*, *he1a*, *sox32*, *sox17* and *pax1b* on force-directed layout of prechordal plate and anterior endoderm cells constructed by Palantir. (G-H) Force-directed layout of hatching gland and pharyngeal pouch (anterior endoderm) cells of zebrafish embryos from blastula stage to organogenesis stage[3]. Cells are colored by Palantir[4] pseudotime, differentiation potential, branch probabilities of hatching gland (PP) and pharyngeal pouch (endo.phary) (H) and the expression levels of key marker genes (G). Yellow dotted frame indicates the progenitors. (I) Branch probabilities of five cells randomly selected from progenitor cells. Bars are colored by cell types as the legend shown on the top left of the panel.

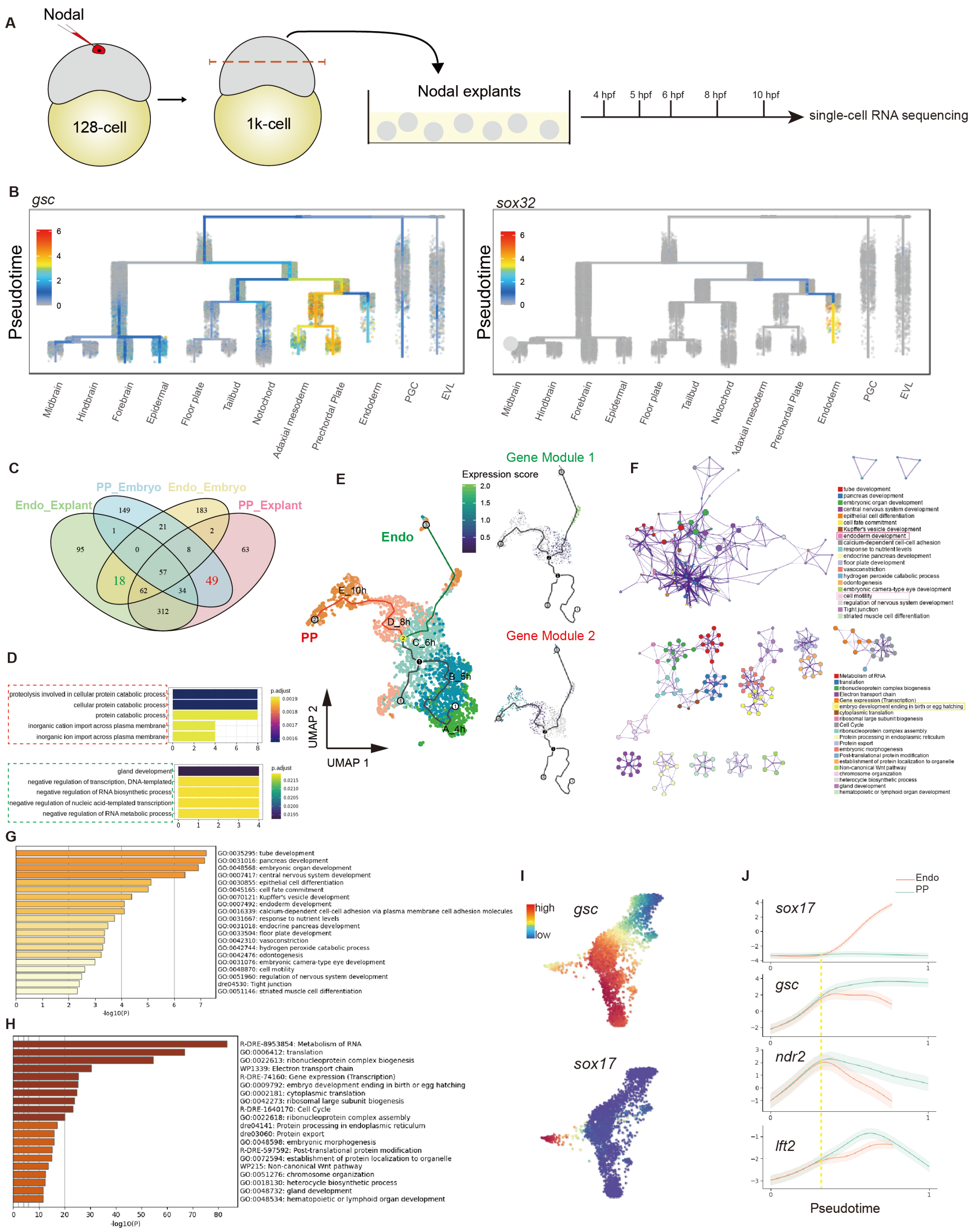


**Figure S2 Investigating the molecular mechanisms of prechordal plate and anterior endoderm separation by Nodal explant single-cell datasets from blastula stage to the end of gastrulation.** (A) Schematic diagram showing the experimental workflow of single-cell RNA-seq for Nodal explants from 4 hpf to 10 hpf. (B) URD plots showing the expression of *gsc* and *sox32* on the single-cell trajectory constructed by single-cell datasets of Nodal explants. Cells are colored by the expression levels. (C) Venn plot showing the numbers of the shared highly expressed genes in the prechordal plate fate of embryos (blue) and Nodal explants (pink)

and in the anterior Endo fate of embryos (yellow) and Nodal explants (green). The gene numbers are showed in the plot accordingly. 18 conserved PP fate genes (green) and 49 anterior Endo fate genes (red) are identified, indicating these genes are conserved between embryos and Nodal explants. (D) Bar plots showing the gene set enrichment analysis using gene sets from (C) with the top 5 GO enriched terms shown. Red dotted frame indicates the enriched GO terms using 18 conserved PP fate genes, and green dotted frame indicates the enriched GO terms using 49 conserved anterior Endo fate genes. The bar is colored by adjusted p-value, and the length of the bar indicates the gene number of each GO term. (E) The single-cell trajectory of prechordal plate and anterior endoderm analyzed by Monocle 3 and visualized by UMAP. Cells are colored by developmental stages (left), and the expression levels of gene module 1 (top right) and gene module 2 (bottom right). Gene module 1 and gene module 2 indicate the co-upregulated genes in anterior Endo fate and in PP fate respectively. (F) Network layout of the enriched GO terms of gene module 1 (top) or gene module 2 (bottom). The node size is scaled by the number of input genes belonging to that term, and the color of nodes represents its cluster identity which is labeled in the right. Terms are linked by an edge (similarity score > 0.3, the thickness of the edge indicates the similarity score). (G-H) Bar plots showing the enriched GO terms achieved through the genes from gene module 1 (G) or gene module 2 (H). (I) Force-directed layout of prechordal plate and anterior endoderm cells of Nodal explants. Cells are colored by the expression levels of *gsc* (top) and *sox17* (bottom). (J) Gene expression trends of anterior Endo (*sox17*) and prechordal plate (*gsc*) marker and Nodal direct targets (*ndr2* and *lft2*) along these two differentiation cell trajectories. The yellow dotted line indicates the point of *sox17* start to be differentially expressed in these two cell trajectories.

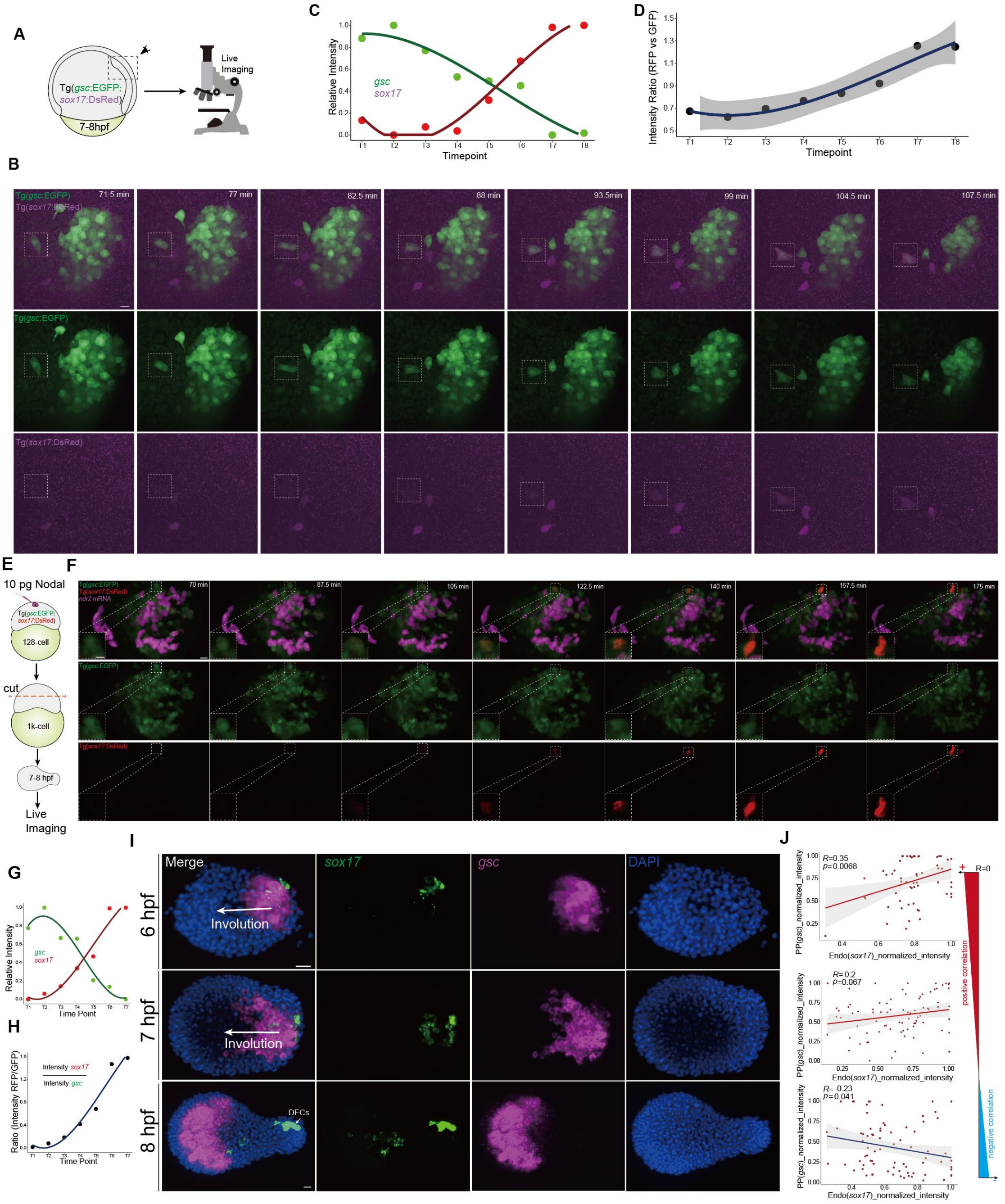


**Figure S3 Live imaging analysis showing transitions from prechordal plate progenitor to anterior Endo in Tg(*gsc*:EGFP;*sox17*:DsRed) transgenic embryos.** (A) Schematic diagram showing the workflow for live imaging analysis. A representative Tg(*gsc*:EGFP;*sox17*:DsRed) transgenic embryo was imaged at 7-8 hpf. The focused region was indicated by black dotted frame. (B) Time series of 3D reconstruction of a representative embryo. (C) Scatter plot and the inferred linear regression showing the temporal profiles of the intensities of *sox17* (RFP) & *gsc* (GFP) in the highlighted cell from (B). (D) Scatter plot and the inferred linear regression indicating the temporal profiles of the ratio between RFP intensity and GFP intensity. Each live imaging experiment was performed on at least 3 independent replicates. (E) Schematic diagram indicating the live imaging analysis for Nodal injected explants which were generated from Tg(*gsc*:EGFP;*sox17*:DsRed) embryos. (F) Time series of 3D reconstruction of a representative Nodal injected explant showed in (E). The cells that were labeled with magenta represented the descendants of the Nodal injected (fluorescent dye plus *ndr2* mRNA) cells at 128-cell stage. Images on the bottom left corner showed enlarged views of the white dashed box regions. (G-H) Temporal profiles of *sox17* (RFP) & *gsc* (GFP) (G) and the ratio between RFP intensity and GFP intensity (H) in the highlighted cell from (F). The intensity of RFP/GFP channel increases/decreases during development. (I) FISH of *sox17* and *gsc* in Nodal explants (n = 5/5, 6 hpf; n = 4/6, 7 hpf; n = 5/7, 8 hpf; n: embryos were imaged, expression observed/total imaged). (J) Scatter plots and inferred linear regression comparing the relative intensities of *sox17* and *gsc* in Nodal explants at 6 hpf (top), 7 hpf (middle) and 8 hpf (bottom). Each FISH experiment was performed for at least 3 independent replicates (technical replicates), more than 30 embryos were analyzed. Scale bar: 20 μm (F and I), 15 μm (B) and 10 μm (subpanel of F).

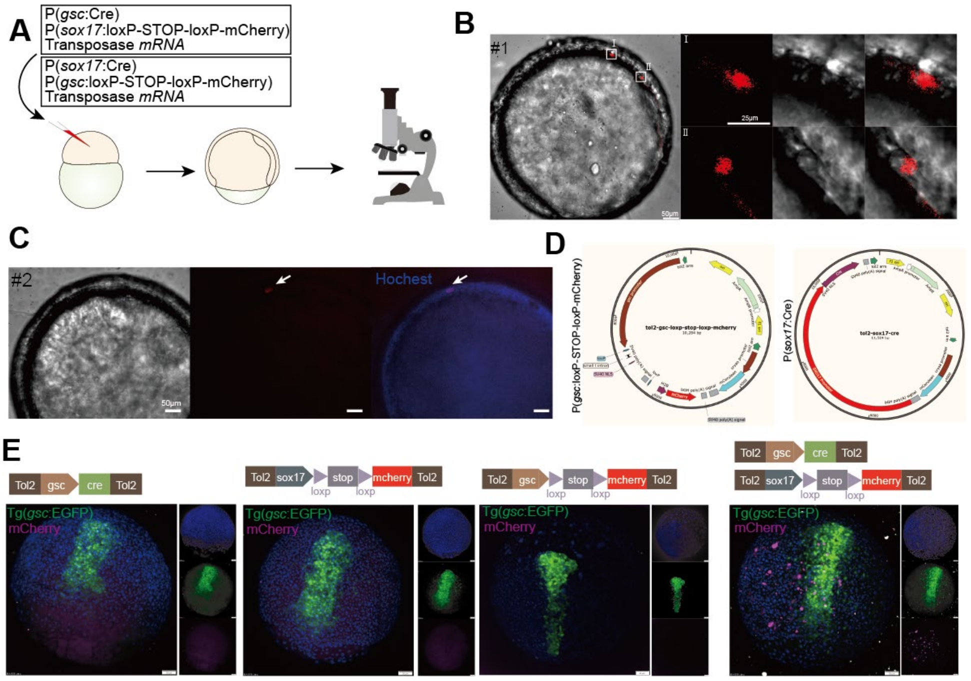


**Figure S4 Inserting *sox17*:Cre and *gsc*:loxP-STOP-loxP-mCherry cassettes in zebrafish embryos.** To support the claim that anterior endoderm originates from prechordal plate progenitor cells, a mosaic transgenic line was generated by inserting the *sox17*:Cre and *gsc*:loxP-STOP-loxP-mCherry cassettes or *gsc*:Cre and *sox17*:loxP-STOP-loxP-mCherry cassettes. This experimental setup allows for capturing the transition from prechordal plate progenitor cells to anterior endoderm. (A) Schematic diagram showing the workflow for generating the mosaic transgenic line containing *sox17*:Cre and *gsc*:loxP-STOP-loxP-mCherry cassette. The plasmids P(*sox17*:Cre), P(*gsc*:loxP-STOP-loxP-mCherry) and transposase mRNA were co-injected at 1-cell stage. (B) Merged images indicating that *sox17* positive cells (endoderm) were derived from *gsc* positive cells (prechordal plate). The right panels show the enlarged view of the regions in the left panel. Two transformed cells were observed in #1 embryo. (C) One transformed cell was also observed in #2 embryo. (D) The plasmid structure of P(*sox17*:Cre) and P(*gsc*:loxP-STOP-loxP-mCherry). (E) Fluorescent images of endodermal cells (mCherry-positive) in the Tg(*gsc*:EGFP) transgenic lines following injection of the indicated plasmids. The experiment was performed on at least 3 independent replicates. Scale bar: 50 μm (B, C and E) and 25 μm (right panel of B).

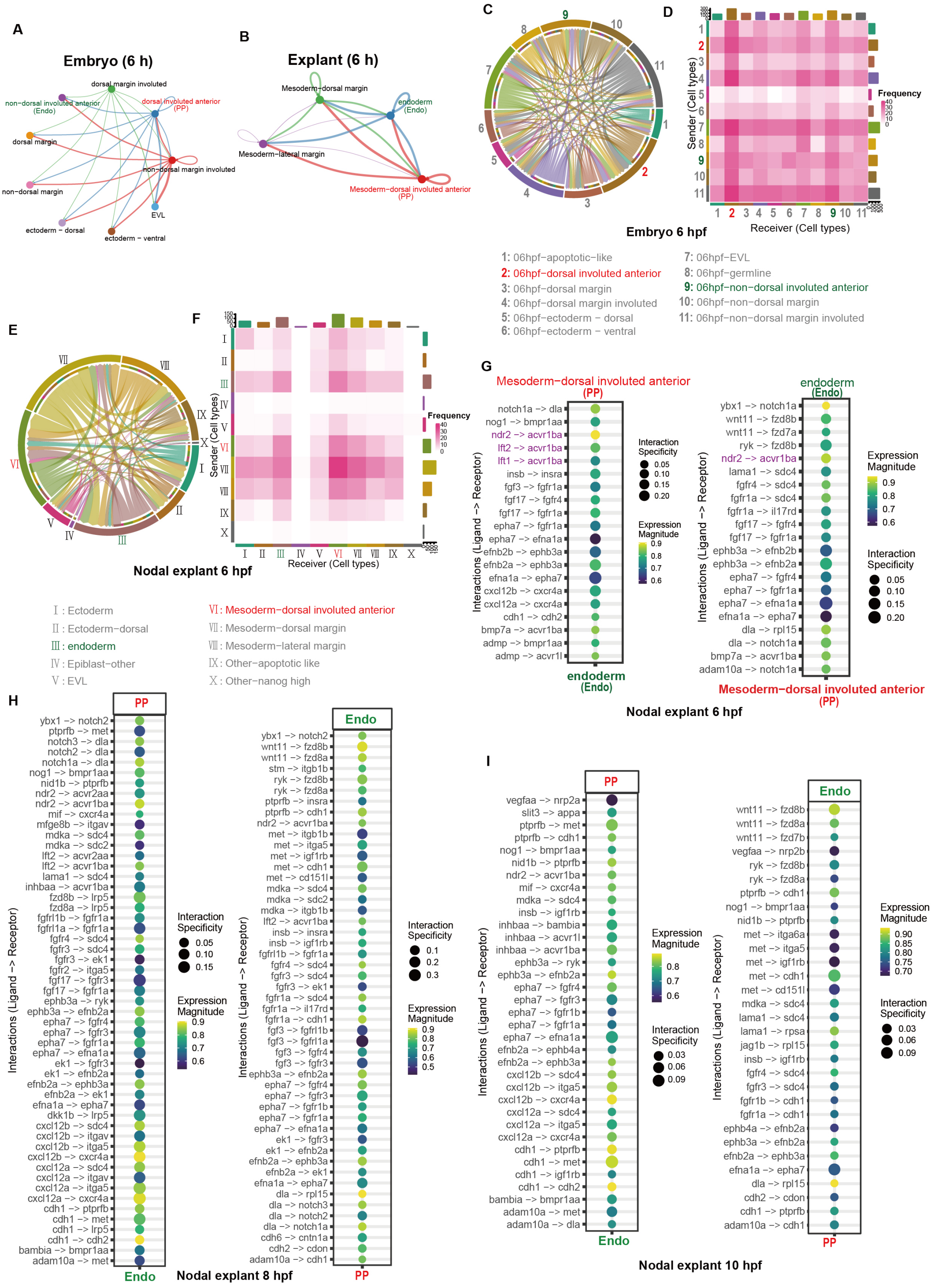


**Figure S5 Analysis of ligand-receptor interactions among different cell types in embryos and Nodal explants.** (A-B) Circle plot showing the Nodal signaling pathway network among the cell types in wild-type embryos (A) and Nodal explants (B) at 6 hpf. The thickness of the edges is proportional to the strength of the interactions. (C-F) Frequency chord diagrams and frequency heatmaps visualizing the frequencies of interactions for each pair of cell types in zebrafish embryos (C and D) and Nodal explants (E and F) at 6 hpf. (G-I) Dot plots visualizing ligand-receptor interactions between PP and anterior Endo in 6 hpf (G), 8 hpf (H) and 10 hpf Nodal explants (I). The interactions from PP (ligand) to anterior Endo (receptor) and from anterior Endo (ligand) to PP (receptor) are plotted respectively on the left and right (G, H and I). The dot size indicates the specificity of the interactions between the ligand and receptor, and the color scale represents the magnitude of expression of the ligand and receptor. Interactions are filtered by an aggregate rank < 0.05.

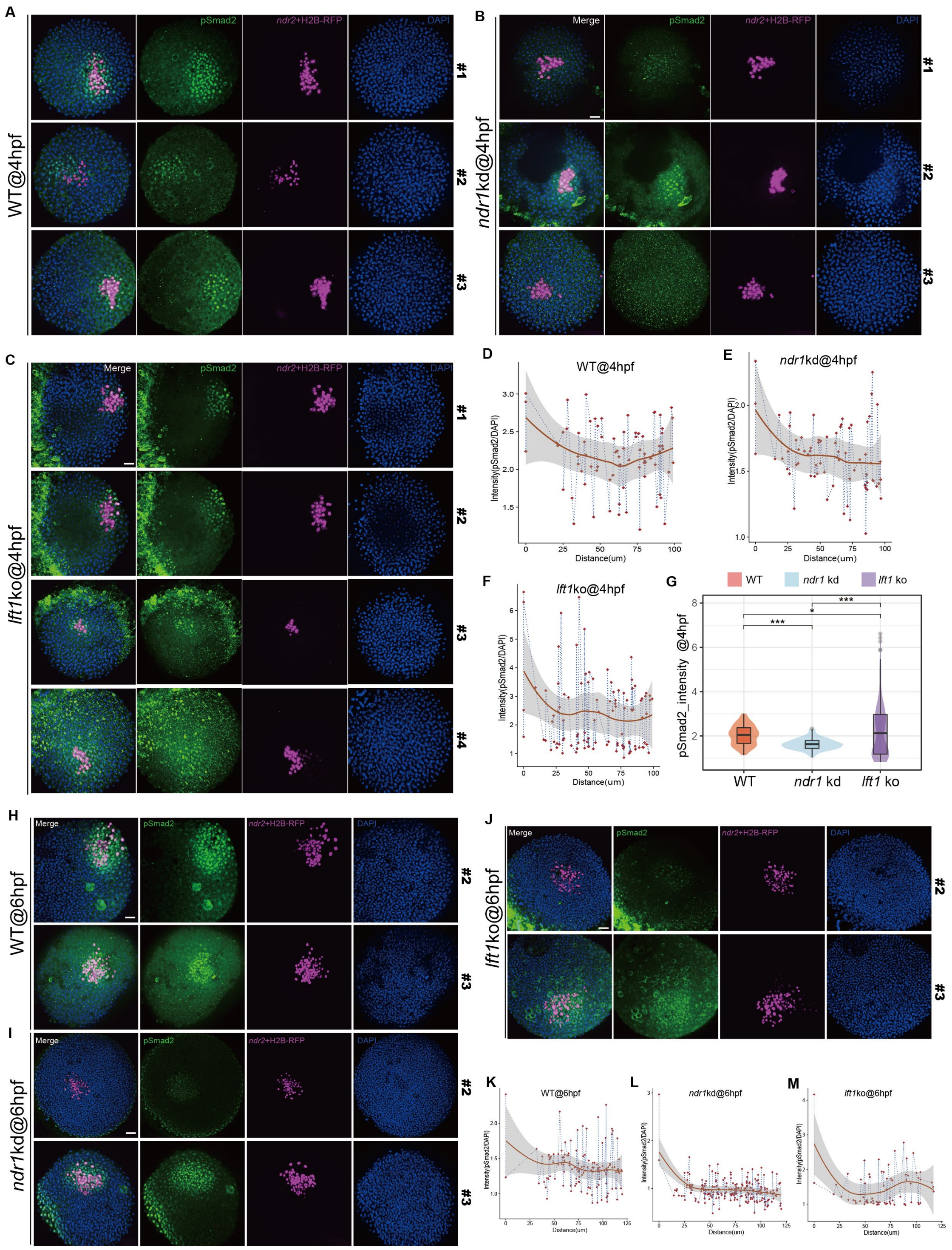


**Figure S6 Analyzing Nodal signaling activity by pSmad2 immunostaining.** (A-F) 3D reconstructed confocal scanned images showing pSmad2 levels in zebrafish embryonic animal pole stimulated by Nodal injection at 128-cell stage. pSmad2 immunostaining (A-C) and its quantifications (D-F) in wild-type embryos (A and D), *ndr1* morphants (B and E) and *lft1* mutants (C and F) at 4 hpf. (G) Volin plot showing comparisons of pSmad2 levels in wild-type embryos, *ndr1* morphants and *lft1* mutants at 4 hpf (D-F). Statistical differences between two samples were evaluated by Student's t-test. *indicates P-value < 0.05, **indicates P-value < 0.01 and ***indicates P-value < 0.001. (H-M) pSmad2 immunostaining (H-J) and its quantifications (K-M) in wild-type embryos (H and K), *ndr1* morphants (I and L) and *lft1* mutants (J and M) at 6 hpf. Each immunostaining experiment was performed for at least 3 independent replicates (technical replicates). Scale bar: 50 µm (A-C, H-J).

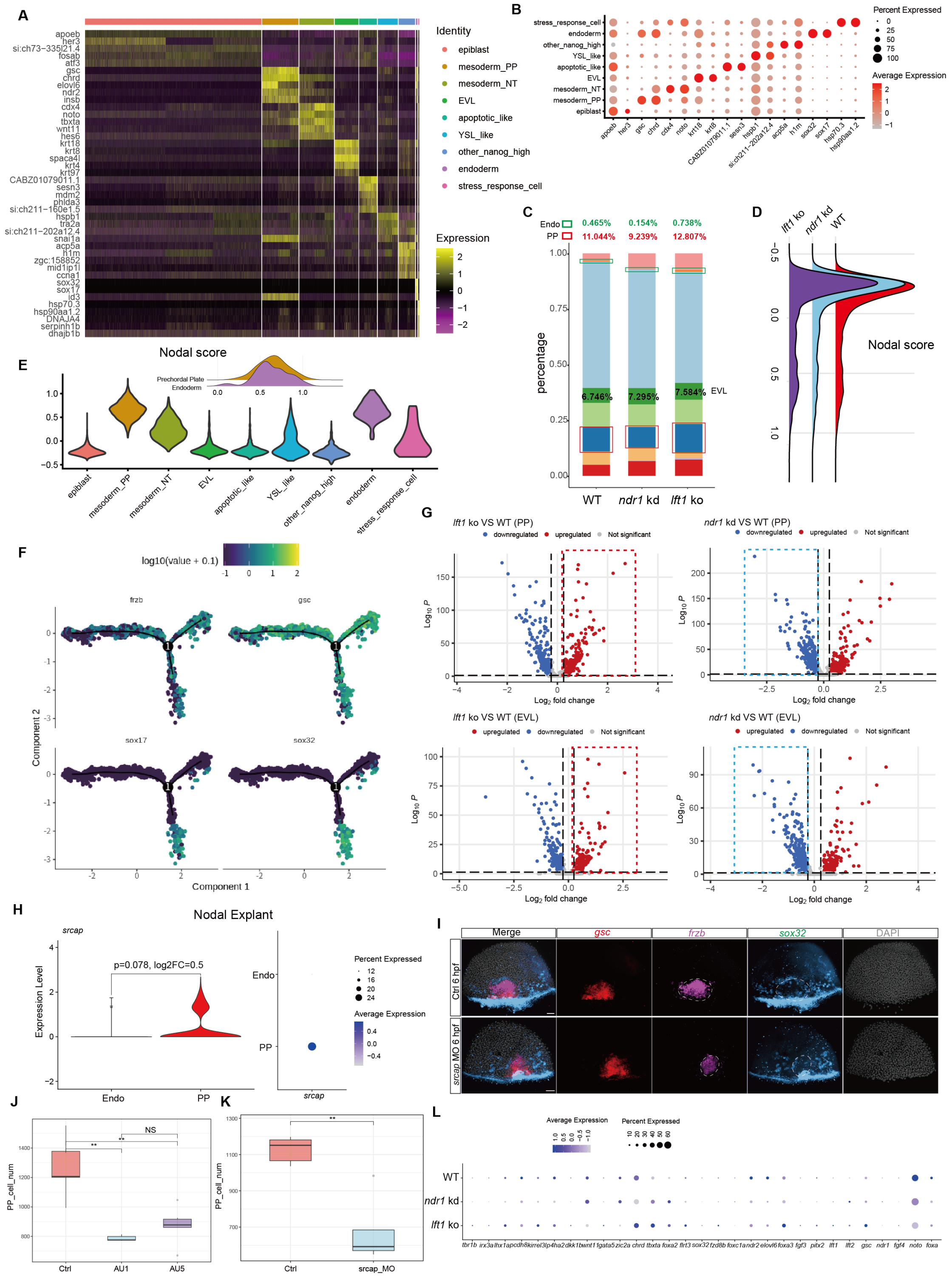


**Figure S7 Single-cell RNA sequencing of wild-type, *ndr1*-morphant and *lft1*-mutant Nodal explants.** (A) Heatmap showing the expression of the top 5 enriched markers in each cell type from integrated single-cell datasets of wild-type, *ndr1*-morphant and *lft1*-mutant Nodal explants. (B) Dot plot showing the expression of the top 2 enriched markers in each cell type of the integrated single-cell datasets. (C) Stacked bar plots showing the proportions of each cell type in wild-type, *ndr1*-morphant and *lft1*-mutant Nodal explants. The proportion values of

PP, anterior Endo and EVL are shown. (D) Ridge plot showing the Nodal score in wild-type, *ndr1*-morphant and *lft1*-mutant Nodal explants. (E) Violin plot showing the Nodal score in each cell type of the integrated single-cell datasets. (F) The expression of *frzb*, *gsc*, *sox17* and *sox32* on the pseudotime trajectory tree of prechordal plate and anterior endoderm from the integrated single-cell datasets of wild-type (left), *ndr1*-morphant (middle) and *lft1*-mutant (right) Nodal explants. (G) Volcano plots of DE genes in each comparison highlighting upregulated genes in red and downregulated genes in blue. DE genes are recognized by $\log_2 FC > \pm 0.25$ (the black vertical lines) and a Bonferroni-adjusted p-value < 0.05 (the black horizontal line). (H) Violin plot (left) and Dot plot (right) indicating the expression levels of *srcap* in PP and anterior Endo cells in Nodal explants. (I) HCR co-staining of *gsc*, *frzb*, and *sox32* in wild-type embryos and embryos with *srcap* knockdown. Embryos at 6 hpf (Ctrl, n = 5/5; *srcap* MO, n = 4/5; n: embryos were imaged, expression observed/total imaged) were evaluated, more than 40 embryos were analyzed. (J and K) Box plot showing the cell numbers of PP in Figure 4H (J) and Figure 4L (K). Statistical differences between two samples were evaluated by Student's t-test (I and M). *indicates P-value < 0.05; NS indicates P-value >=0.05. (L) Dot plot showing the expression levels of Nodal target genes used to calculate the Nodal activity score across WT, *ndr1* knockdown, and *lft1* knockout samples from single-cell RNA-seq data of Nodal explants. Scale bar: 50 μm (I).

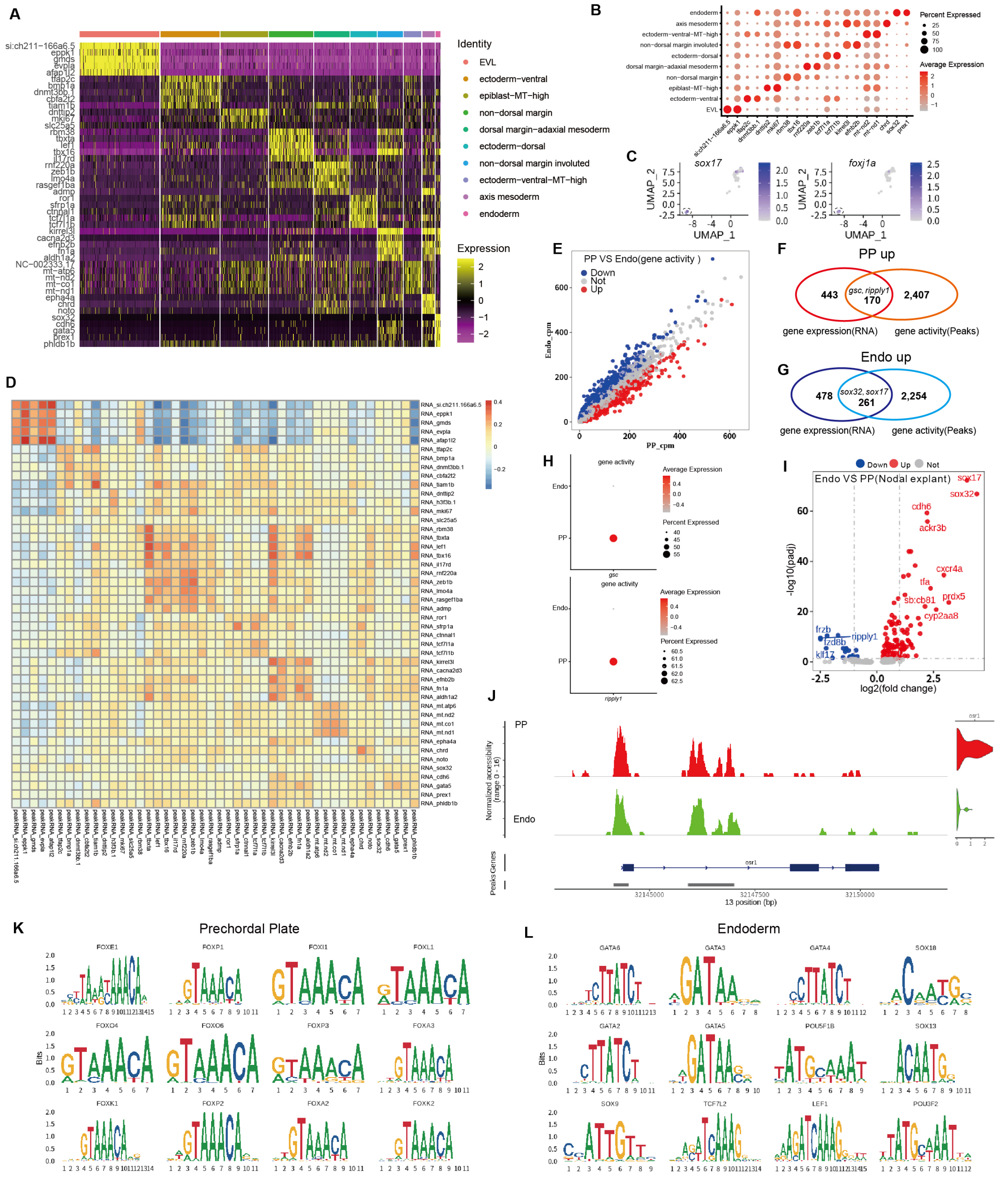


**Figure S8 Single-cell multiomics of 6 hpf zebrafish embryos.** (A) Heatmap showing the expression (RNA level) of the top 5 enriched markers in each cell type from single-cell multiomics datasets of 6 hpf zebrafish embryos. (B) Dot plot showing the expression of the top 2 enriched markers in each cell type of the single-cell multiomics datasets. (C) The expression of *sox17* and *foxj1a* on the UMAP of prechordal plate cells, endoderm cells and DFCs. (D) Heatmap showing correlation analysis of marker genes in each cell type between the RNA expression levels of scRNA-seq data and the relative RNA-peak levels of single-cell ATAC data. The correlations are measured by Spearman's tests, and the intensity of color scale represents correlation coefficients with high in red and low in blue. (E) Scatter plot showing gene activity levels (calculated from chromatin openness of each gene, **see Methods**) in prechordal plate and endoderm cells. Red dots and blue dots indicate up-regulated genes and down-regulated genes (identified by differential expression analysis) in prechordal plate respectively. (F-G) Venn plots showing the overlapped up-regulated genes at gene expression levels and gene activity levels in prechordal plate cells (F) and endoderm cells (G). (H) Dot plots showing the gene activity levels of *gsc* and *ripply1* in prechordal plate and endoderm. (I) Volcano plot showing the differentially expressed genes in prechordal plate and endoderm of Nodal explants at 6 hpf. (J) Track plot showing chromatin accessibility of each cell type on the gene locus of *osr1*. Violin plot showing the RNA levels of *osr1* in prechordal plate and endoderm (on the right of each panel). (K and L) Motif plots showing the enriched motifs and related genes in prechordal plate (K) and endoderm (L).

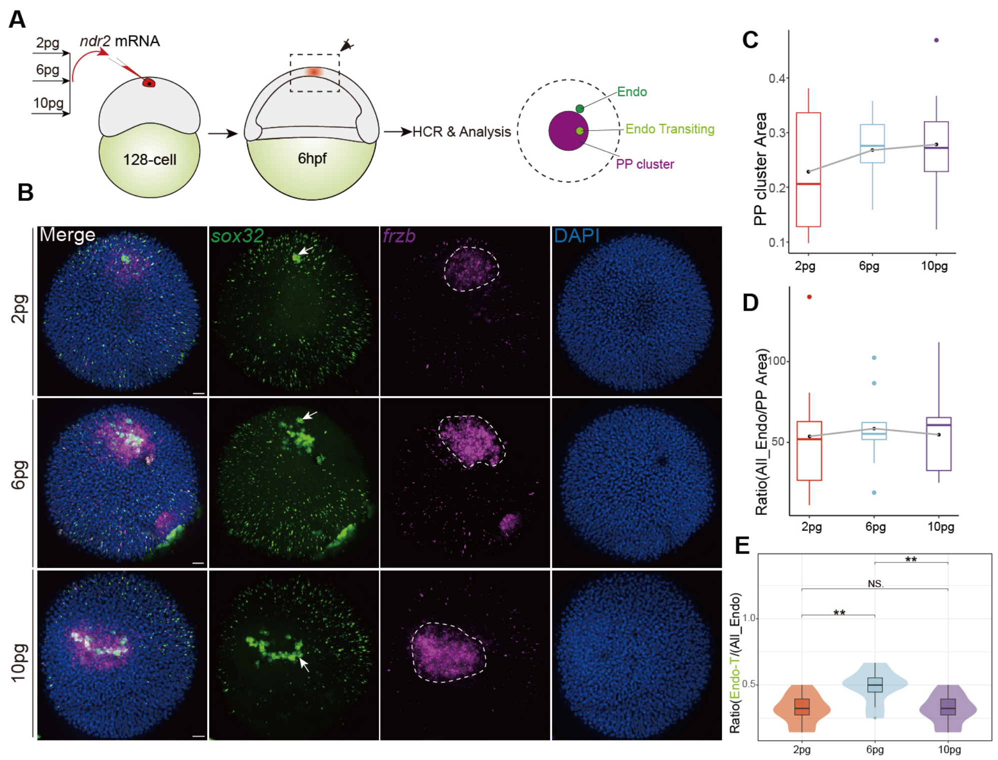


**Figure S9 The involvement of Nodal concentration in PP and anterior Endo separation.** (A) Schematic diagram illustrating the strategy employed to investigate the influence of Nodal concentration on the cell fate separation of PP and anterior Endo. Different dosages of Nodal mRNA were injected into one zebrafish animal pole blastomere at the 128-cell stage. The effects on mesendoderm cell fate separation were assessed by quantifying the area of the PP cluster and the number of anterior Endo cells. (B) HCR co-staining of *sox32* and *frzb* on the embryos injected with different dosages of Nodal mRNA (n = 7/10, 2 pg; n = 10/11, 6 pg; n = 11/13, 10 pg; n: embryos were imaged, expression observed/total imaged). (C) Box plot quantifying the area of PP cell clusters in (B). (D) Box plot showing the ratio of anterior Endo cell number to the area of PP cell clusters. (E) Volin plot showing the ratio of transiting anterior Endo cell number to the total anterior Endo cell number. Each HCR was performed for at least 2-3 independent replicates (technical replicates), more than 20 embryos were analyzed. Scale bar: 30 μm (B).

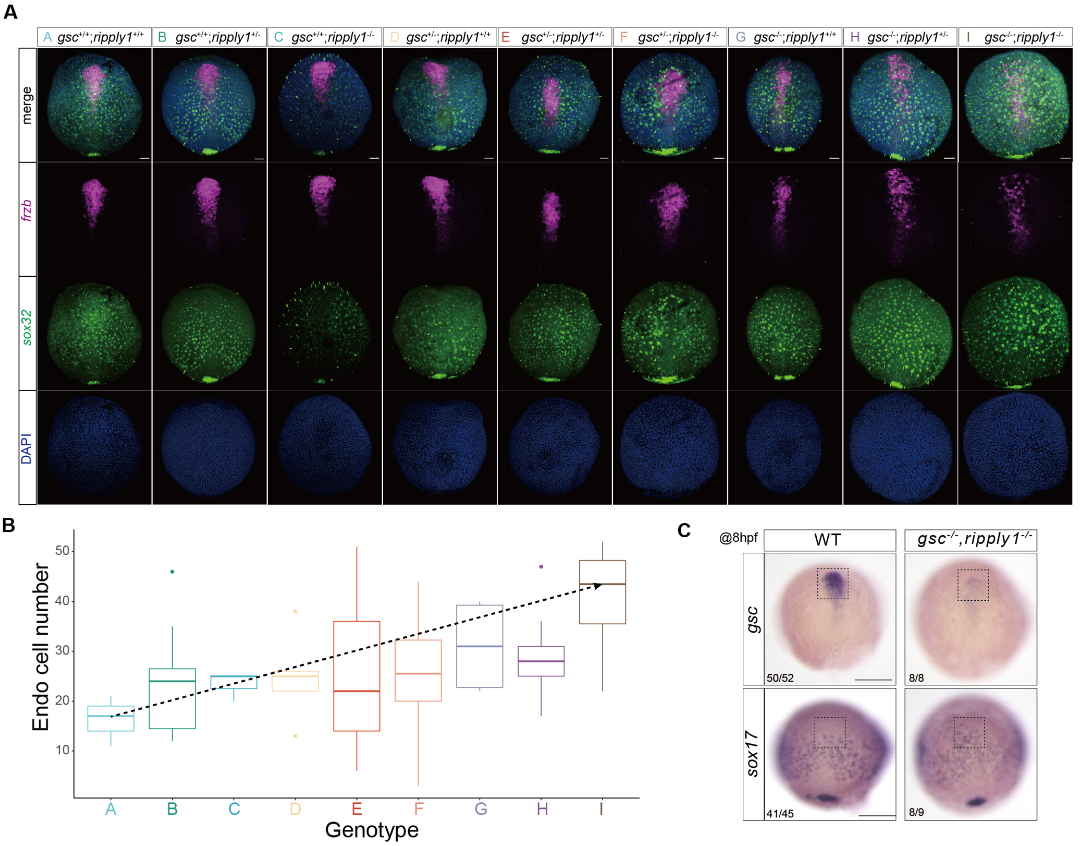


**Figure S10 Investigation of the cell fate specification of prechordal plate and anterior endoderm in the descendant embryos of Tg(*gsc*$^{+/-}$;*ripply1*$^{+/-}$).** (A) HCR co-staining of *sox32* and *frzb* in the descendant embryos of Tg(*gsc*$^{+/-}$;*ripply1*$^{+/-}$) at 8 hpf. Here embryos of nine different genotypes are shown [Tg(*gsc*$^{+/+}$;*ripply1*$^{+/+}$), n = 4/5; Tg(*gsc*$^{+/+}$;*ripply1*$^{+/-}$), n = 11/13; Tg(*gsc*$^{+/+}$;*ripply1*$^{-/-}$), n = 4/6; Tg(*gsc*$^{+/-}$;*ripply1*$^{+/+}$), n = 5/6; Tg(*gsc*$^{+/-}$;*ripply1*$^{+/-}$), n = 12/15;

Tg($gsc^{+/-}$;$ripply1^{-/-}$), n = 10/12; Tg($gsc^{-/-}$;$ripply1^{+/+}$), n = 4/5; Tg($gsc^{-/-}$;$ripply1^{+/-}$), n = 9/12; Tg($gsc^{-/-}$;$ripply1^{-/-}$), n = 5/5; expression observed/total imaged]. (B) Boxplot showing the quantification of the numbers of anterior Endo cells identified in (A). The black dashed line represents the increasing trend in the number of anterior Endo cells with the increased number of mutant alleles of *gsc* and *ripply1*. (C) ISH of *gsc* (up) and *sox17* (bottom) in WT (left) and Tg($gsc^{-/-}$;$ripply1^{-/-}$) (right) embryos at 8 hpf. Each HCR and ISH experiment was performed for at least 3 independent replicates (technical replicates). Scale bar: 50 μm (A) and 200 μm (C).

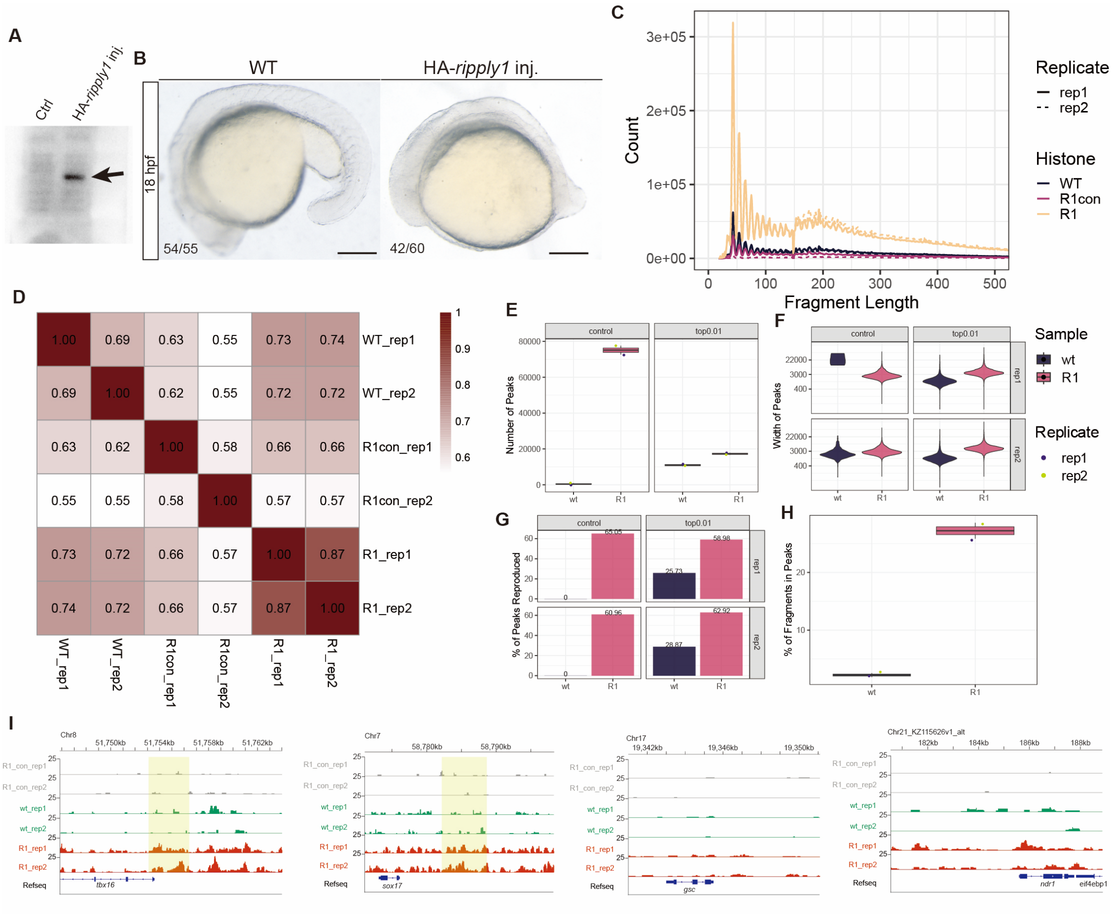


**Figure S11 Exploration of Ripply1 downstream targets by CUT&Tag experiment.** (A) Anti-HA western blot showing the expression of HA-Ripply1 fusion protein in HA-*ripply1*-injected embryos. (B) The phenotype of 18 hpf embryos after HA-*ripply1* mRNA injection. Overexpression of HA-*ripply1* led to dorsalized embryos, which was consistent with previous findings that *ripply1* partially functioned as dorsal organizer[5]. (C) Plot of fragment lengths with single-basepair resolution. (D) Correlation heatmap showing the reproducibility between replicates and across conditions. Numbers indicate the Pearson correlation coefficient. (E, F, G,

H) Assessment of peaks called by SEACR, including the peak number (E), peak width (F), peak reproducibility (G) and the fraction of reads in peaks (H). Control indicates the peaks are called using technical control as control; while top0.01 indicates the top 1% of enrich regions are selected as peaks by AUC. (I) Screenshot of R1_con (technical control, without primary antibody), WT (ddH2O-injected) and R1 (HA-ripply1-injected) CUT&Tag peaks around *tbx16*, *sox17*, *gsc* and *ndr1* loci. Scale bar: 200 μm (B).

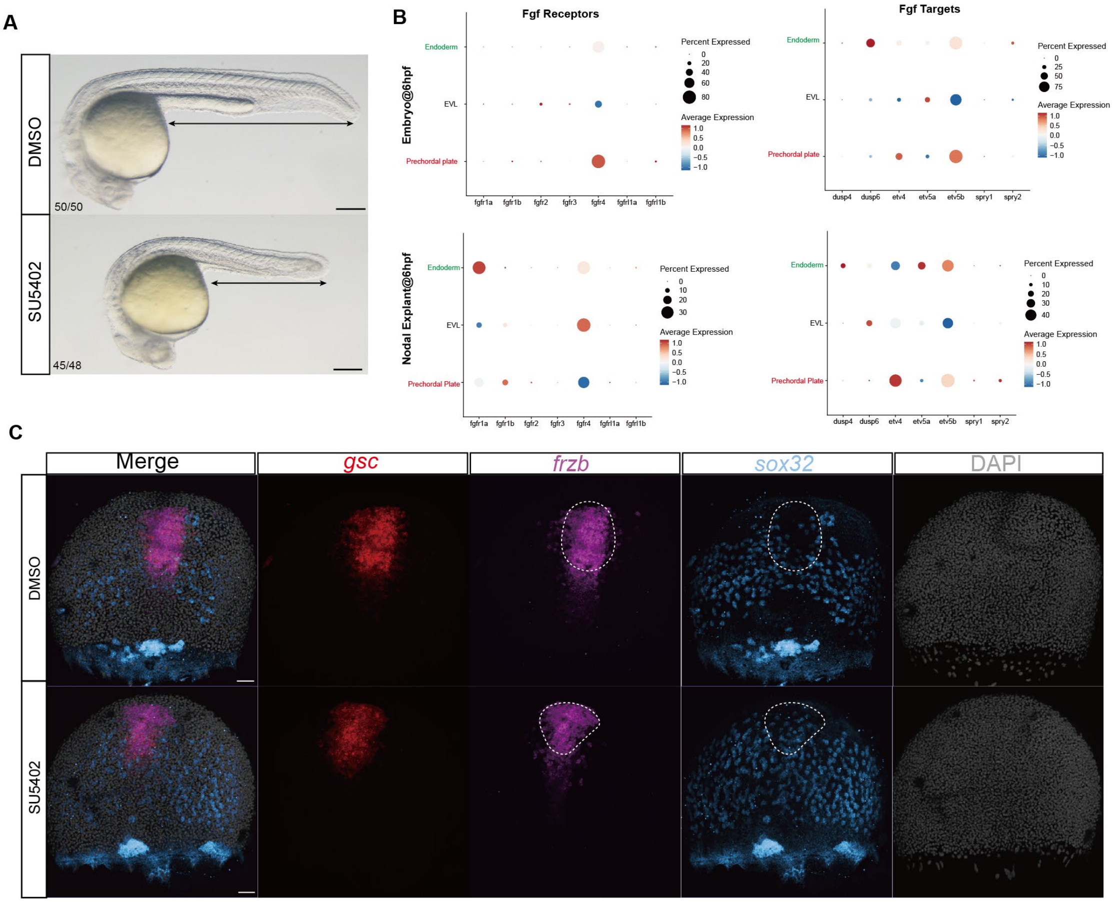


**Figure S12 Investigating the role of Fgf signaling in the regulation of cell fate differentiation between prechordal plate and anterior endoderm.** (A) Bright filed images showing that inhibiting Fgf signaling by SU5402 results in the developmental defect characterized by a shortened tail, which aligns with the findings from a previous study[6]. (B) Dot plots showing the high expression levels of some Fgf receptors and downstream target genes of Fgf signaling in both PP and anterior Endo cells observed in embryo (top) and Nodal explant (bottom) single-cell data. (C) HCR co-staining of *gsc*, *frzb* and *sox32* in wild-type and SU5402-treated embryos (DMSO, n = 7/7; SU5402, n = 4/5; expression observed/total imaged).

The samples were examined at 8 hpf, with the region of anterior mesendoderm indicated by white dotted lines. The experiment was replicated for at least 3 times. Each HCR experiment was performed for at least 3 independent replicates (technical replicates). Scale bar: 50 μm (C) and 200 μm (A).

**Movie S1. Time-lapse imaging of Nodal explants.** Time-lapse imaging of a representative Nodal explant generated from a Tg(*gsc*:EGFP;*sox17*:DsRed) embryo from 7 hpf onwards (n = 10, N = 5). Cells marked by magenta indicate the descendants of Nodal injected clones. Prechordal plate progenitors are marked by the expression of EGFP, and anterior endoderm cells are marked by the expression of DsRed. "n" indicates the number of imaged embryos, and "N" indicates the number of replicated experiments. 20X objective lens was used.

**Movie S2. Time-lapse imaging of Tg(*gsc*:EGFP;*sox17*:DsRed) transgenic embryos.** A representative Tg(*gsc*:EGFP;*sox17*:DsRed) embryo was taken for time-lapse imaging from 7 hpf onwards (n = 14, N = 8). Prechordal plate progenitors are marked by the expression of EGFP, and anterior endoderm cells are marked by the expression of DsRed. "n" indicates the number of imaged embryos, and "N" indicates the number of replicated experiments. 20X objective lens was used.

**Movie S3. Time-lapse imaging of Tg(*gsc*:EGFP;*sox17*:DsRed) transgenic embryos using a higher magnification objective lens.** A representative Tg(*gsc*:EGFP;*sox17*:DsRed) embryo was taken for time-lapse imaging from 8 hpf onwards (n = 8, N = 5). 40X oil immersion lens was used.

**Movie S4. Time-lapse imaging of Tg(*gsc*:EGFP;*sox17*:DsRed) transgenic embryos.** A representative Tg(*gsc*:EGFP;*sox17*:DsRed) embryo was taken for time-lapse imaging from 8 hpf onwards (n = 8, N = 5). 40X oil immersion lens was used. Imaris was used to track the cells that transiting from EGFP positive cells to RFP positive cells.

**Movie S5. Time-lapse imaging of Tg(*gsc*:EGFP;*sox17*:DsRed) transgenic embryos.** A representative Tg(*gsc*:EGFP;*sox17*:DsRed) embryo was taken for time-lapse imaging from 6 hpf onwards (n = 10, N = 7).

**Table S1. Primers used in this study.**

**Data S1. Conserved genes that highly expressed in anterior endoderm or prechordal plate fate (embryos and Nodal explants).** These trajectory-specific highly expressed genes were identified by URD trees constructed from embryos and Nodal explants, and the overlapped genes were recognized as conserved genes.

**Data S2. Marker genes of integrated single-cell RNA-seq data of Nodal explants (wild-type, *lft1* mutants and *ndr1* morphants) at 6 hpf.**

**Data S3. Enriched GO terms identified by gene sets that highly expressed in the progenitor cells of prechordal plate and anterior endoderm in the integrated single-cell RNA-seq data of Nodal explants.**

**Data S4. Enriched GO terms identified by gene sets that highly expressed in the prechordal plate cells in the integrated single-cell RNA-seq data of Nodal explants.**

**Data S5. Enriched GO terms identified by gene sets that highly expressed in the anterior endoderm cells in the integrated single-cell RNA-seq data of Nodal explants.**

**Data S6. Marker genes of single-cell multiomics data of zebrafish embryos at 6 hpf.** The genes were identified by analyzing the RNA levels of the single-cell data.

**Data S7. Differentially expressed genes in prechordal plate and endoderm (prechordal plate versus endoderm) identified by comparing the RNA levels between these two cell types.**

**Data S8. Differentially expressed genes in prechordal plate and endoderm (prechordal plate versus endoderm) identified by comparing the gene activation levels (peak intensity) between these two cell types.**
