## Supplementary material for "Ripply1 and Gsc collectively suppress anterior endoderm differentiation from prechordal plate progenitors": TableS1

Table S1 Primers used in this study.

| genes | forward primers | reverse primers |
| --- | --- | --- |
| PCR Primers and primers for sequencing | | |
| *gsc* | 5’-GTATAAAGCAGAGCAGGAGG-3’ | 5’-CTGGTGTATGCGTTCTTGCG-3’ |
| *ripply1* | 5’-ACAGGGATCATTGTCGTCTC-3’ | 5’-GTAGGCTACTCACACAAGCC-3’ |
| *gsc* sequencing |  | 5’-GAATACACGGACACTGTTGCGAA-3’ |
| *ripply1* sequencing | 5’-GTGCTTTGCCACTCCTCATACTA-3’ |  |
| Primers for constructing HA-*ripply1* plasmid | | |
| HA-*ripply1*  F1 | 5‘-tcaaggcctctcgagcctctagaATGTACCC  ATACGATGTTCCAGATTACGCT-3’ | 5’-TACGACTCACTATAGTTCTAGAtca  gttgaaagctgtgaagtga-3’ |
| HA-*ripply1*  F2 | 5’-ACGATGTTCCAGATTACGCTGGCAG  CATGAATTCTGTGTGCTTTGCCACT-3’ |  |
| qPCR primers | | |
| *GSC* | 5’-TCAACCAGCTGCACTGTCGGC-3’ | 5’-TCCATTTGGCGCGGCGGTTC-3’ |
| *SOX17* | 5’-TGAACGCTTTCATGGTGTGGGC TAAGGACGAG-3’ | 5’-CGGTACTTGTAGTTGGGGTGGTCC  TGCATGTG-3’ |
| *srcap* | 5’-CTCACTCCCATTGAGCGTTATGC CA-3’ | 5’-CGAGGGCGACGGGATGGAGTAGG CT-3’ |
| *sox10* | 5’- AGCCACAGCCAATCGCATTA-3’ | 5’-AGTCCACTCCGAGAGGCTCC-3’ |
| *18s* | 5’-TCGCTAGTTGGCATCGTTTATG-3’ | 5’-CGGAGGTTCGAAGACGATCA-3’ |
